# Noradrenergic signaling in wakeful states inhibits microglial surveillance and synaptic plasticity in the mouse visual cortex

**DOI:** 10.1101/556480

**Authors:** Rianne D. Stowell, Grayson O. Sipe, Ryan P. Dawes, Hanna N. Batchelor, Katheryn A. Lordy, Jean M. Bidlack, Edward Brown, Mriganka Sur, Ania K. Majewska

## Abstract

Microglia are the innate immune cells of the brain with roles in neuroimmunology and synaptic plasticity. Microglial processes continuously survey the brain parenchyma interacting with synaptic elements and maintaining tissue homeostasis. However, the mechanisms that control surveillance and its role in synaptic plasticity are poorly understood. Microglial dynamics *in vivo* have been primarily studied in anesthetized animals, where slow-wave neural activity resembles sleep-like states. We report that microglial surveillance and injury response in awake animals are reduced compared to when animals are anesthetized, suggesting that arousal state profoundly modulates microglial roles in the physiological brain. Stimulating β_2_-adrenergic receptors recapitulated these observations and also disrupted experience-dependent plasticity only when intact β_2_-adrenergic receptors were present in microglia specifically. These results indicate that microglial roles in surveillance and synaptic plasticity in the healthy brain are modulated by noradrenergic fluctuations between arousal states and raises new considerations for sleep/wake disruption in neurodevelopment and neuropathology.

## Introduction

In recent years, the list of key regulators of synaptic function has expanded to include astrocytes and microglia as critical components in the synaptic unit of the “quad-partite” synapse^1^. However, the extent and mechanisms underlying this microglial role are still under debate. Microglia, the innate immune cells of the central nervous system (CNS), are exquisitely sensitive to changes in brain homeostasis and rapidly alter their morphology to respond to inflammatory signals. In the absence of injury or disease states, microglial arbors are highly ramified with dynamic processes that continually extend and retract through the parenchyma^2,3^. Several recent *in vivo* imaging studies have shown that this microglial process motility permits microglial interactions with neural elements in diverse regions of the CNS^4-7^, allowing microglia to modulate synaptic remodeling and neural plasticity through the release of growth factors and enzymes or through physical synaptic contact^8-10^. While several signaling pathways, such as fractalkine, complement, and purinergic signaling, have been identified as mediating microglial function during neural circuit pruning, development, and plasticity^8,10-13^, it remains largely unknown how other CNS signals, such as neurotransmitters, may regulate microglial function^14^.

Recent evidence suggests that glial physiology can be dramatically altered between sleep and wakeful states. For example, astrocytic calcium signaling increases during wakeful states and is relatively quiescent during sleep-like states^15^. However, studies of microglia have been performed primarily in reduced preparations or in anesthetized animals, leaving open the possibility that microglial dynamics and the resulting interactions with neurons may be altered in awake brains. In addition, extracellular space increases during sleep-like states^16^, which could greatly impact microglial surveillance, and contribute to the enhanced remodeling of synaptic structure^17^ and increased activity-dependent plasticity^18^ that occurs during sleep. Interestingly, in the non-injured brain, microglia have a high and selective expression of the β_2_ adrenergic receptor (β_2_-AR) relative to other CNS cell types^19,20^, suggesting they may have unique responses to norepinephrine (NE), which potently modulates plasticity, learning, attention to salient stimuli, and sensory processing^17^. Furthermore, NE and β_2_-ARs can impact the microglial inflammatory response^19^, IL-1β production^20^, and ATP-driven chemotaxis by inhibiting the P2Y12 Gα_i_ signaling pathway^21^. P2Y12 is also critical for experience-dependent plasticity in the mouse adolescent visual system, and its loss impairs both the microglial response and synaptic plasticity itself^10^ Thus, NE could act through microglial β_2_-ARs to modulate microglial behavior during sleep/wake states and modulate P2Y12-dependent microglial processes that contribute to activity-dependent synaptic rearrangement.

Here, we characterize microglial dynamics in awake mice and describe the effect of microglial β_2_-AR signaling on *in vivo* microglial physiology and functions in synaptic plasticity. We demonstrate that microglia in the awake brain differ from those in anesthetized and “sleep-like” conditions in both baseline morphology and process dynamics as well as injury response. These findings suggest that microglial functions in awake conditions are fundamentally different than what has previously been described in anesthetized animals. We then recapitulate most of these changes through pharmacological stimulation or inhibition of microglial β2-ARs. Finally, we demonstrate that changes in microglial β2-AR and noradrenergic signaling impair ocular dominance plasticity. Our work shows the importance of microglial β_2_-AR signaling in regulating microglial surveillance of the brain, as well as the participation of these cells in experience-dependent plasticity, which provides new consideration for diseases in which sleep or NE homeostasis are disrupted.

## Results

### Anesthesia increases microglial surveillance of the CNS parenchyma

Most studies of microglial dynamics *in vivo* have been conducted in anesthetized animals or in *ex vivo* slice preparations^2,3,10,14^. However, anesthesia substantially alters numerous facets of brain homeostasis through diverse actions on multiple cell types. To determine whether microglial behavior is affected by anesthesia, we imaged microglial dynamics in the same adult animals during anesthetized and wakeful states using the microglia-labeled CX3CR1^GFP^ transgenic mouse line and a chronic cranial window preparation. We found that compared to awake conditions, microglia in mice anesthetized with a commonly used fentanyl cocktail (also containing dexmedetomidine and midazolam) had significantly altered morphology. Quantification of microglial morphology using Sholl analysis revealed that microglia in anesthetized animals had more elaborate process arbors than when the animals were awake (Fig.1a-d; area under the curve (AUC); Paired t-test; p<0.01; Maximum # of intersections Paired t-test; p<0.01). We next examined microglial process coverage across one hour to determine whether the decreased process ramification during wakefulness resulted in decreased parenchymal surveillance. We found a significant reduction in the total parenchymal area surveyed in awake states as compared to anesthetized states, (Fig.1e,f; Paired t-test; p<0.05). To determine whether changes in surveillance were also a result of altered process dynamics, we measured the motility index across one hour. Surprisingly, process motility was higher in awake mice (Fig.1g,h; Supplementary Movies 1 and 2; Paired t-test; p<0.01). A more detailed analysis of the microglial process dynamics indicated that wakefulness did not alter microglial process instability (Fig.1i; Student’s t-test; n.s.), while anesthesia enhanced microglial process stability (Fig.1j; Student’s t-test; p<0.01).

**Figure 1.**
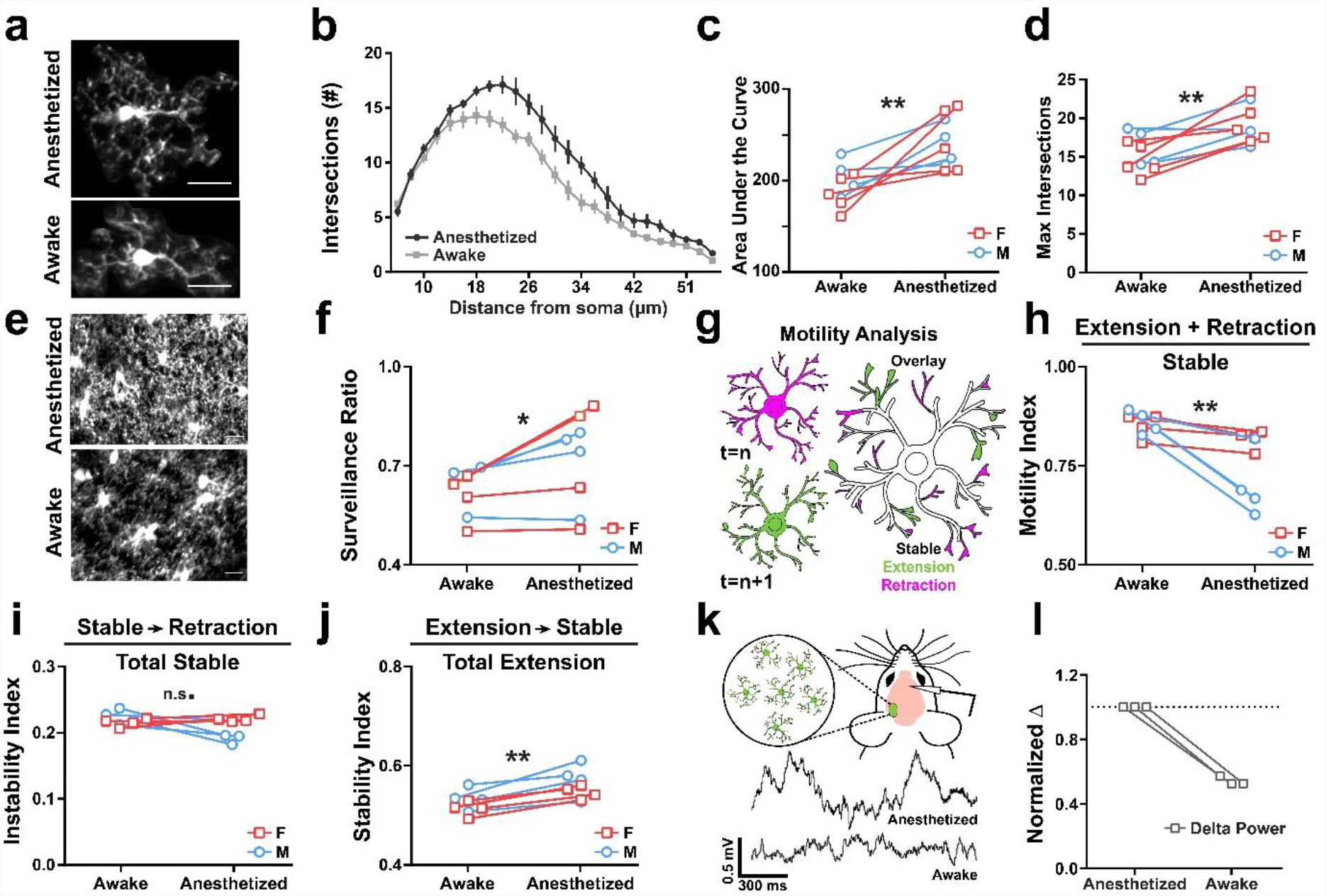
Anesthesia enhances microglial surveillance of the parenchyma. (**a**) Individual microglia from a mouse that was either awake or anesthetized. (**b**) Sholl profile of anesthetized and awake microglia, note a reduction in awake microglial arborization. (**c**) Anesthetized microglia have a greater area under the curve compared to their awake counterparts (n=4m/5f, Paired t-test, P=0.0046, t(8)=3.891). (**d**) Anesthetized microglia have a greater number of maximum process intersections (n=4m/5f, Paired t-test, P=0.0030, t(8)=4.195). (**e**) Max projection of microglial processes over the hour imaging session to observe parenchyma surveillance. (**f**) Microglia in anesthetized animals survey a greater portion of the CNS parenchyma over an hour of imaging (n=4m/4f, Paired t-test, P=0.0087, t(7)=3.606). (**g**) Visual representation of quantification of microglial motility by comparing microglial process movement across consecutive time-points. (**h**) Anesthesia decreases microglial process motility (n=4m/4f Paired t-test, P=0.0120, t(7)=3.366). (**i**) There was no corresponding change in process instability (n=4m/4f, Paired t-test, P=0.2053, n.s., t(7)=1.396). (**j**) Our decreased motility in the anesthetized state did correspond to a significant increase in microglial process stabilization (n=4m/4f, Paired t-test, P=0.0011, t(7)=5.318).(**k**) Example local field potential (LFP) traces from mice in anesthetized and awake mice with thin skull preparations. (**l**) Normalized delta-power from LFP recordings in awake and anesthetized states demonstrating decreased delta-power in the awake state and increased delta power in the anesthetized state. Scale bars 20μm. Graphs show mean ± s.e.m. Individual points represent individual animals.

Despite the animals being habituated to head restraint before imaging, it is possible that changes in microglial morphology and surveillance in awake animals are due to factors released as a result of increased stress. To test this possibility, we imaged microglia in animals that were exposed to restraint stress prior to anesthesia and found no difference in microglial motility, morphology or surveillance (Supplementary Fig.1). We also determined that circadian rhythm did not alter the morphology, surveillance or motility of microglia (Supplementary Fig.2). Based on these findings, we conclude that anesthesia decreases process motility by increasing stability, which results in a more complex morphology and higher process coverage of the brain parenchyma.

Anesthesia has diverse effects on brain function depending on the anesthetic used. To determine the cortical network state under the fentanyl cocktail, we collected local field potential (LFP) recordings in the cortex during anesthesia and wakeful states. As expected, delta wave power was significantly higher under anesthesia (Fig.1k,l), suggesting that the fentanyl cocktail puts the brain in a slow-wave dominated, “sleep-like” state. To disambiguate the effects of the different components of the fentanyl cocktail on microglia, and to more closely simulate a deep sleep state, we repeated our experiments using solely dexmedetomidine (DEX), which is a known sedative that reduces NE release from the locus coeruleus (LC) and approximates natural slow-wave sleep^21^. DEX alone produced a robust increase in the microglial arbor (Fig.2a-d; AUC Paired t-test; p<0.0001; Maximum # of intersections; Paired t-test; p<0.05) and enhanced microglial surveillance (Fig.2e,f; Paired t-test; p<0.05). Microglia in animals dosed with DEX also had a lower rate of process motility recapitulating the fentanyl cocktail results (Fig.2g; Supplementary Video 3; Paired t-test; p<0.001). Likewise, this decreased motility was associated with decreased process instability and increased process stability (Fig.2h,i; instability; Paired t-test; p<0.001; stability; Paired t-test; p<0.001;). These data suggest that inducing a slow-wave sleep like state using DEX is sufficient to elicit a profound enhancement of microglial surveillance of the parenchyma.

**Figure 2.**
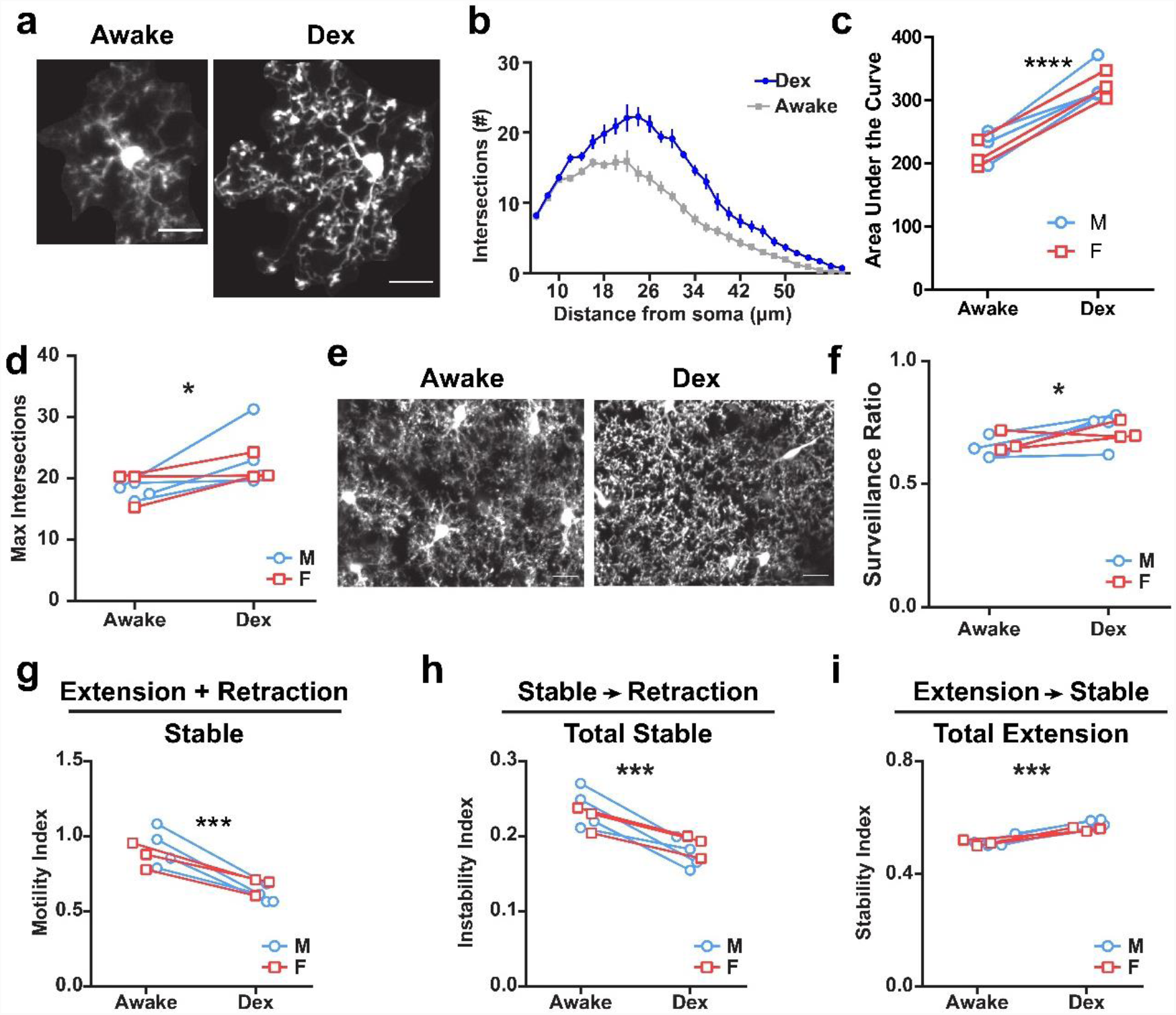
Microglial surveillance is enhanced by inducing a sleep-like state with dexmedetomidine. (**a**) Individual microglia from a mouse that was either awake or anesthetized with dex. (**b**) Sholl profile of dex anesthetized and awake microglia, note the marked enhancement of the microglial arbor with dex. (**c**) Dex anesthetized microglia have a greater area under the curve compared to their awake counterparts (n=4m/3f, Paired t-test, P<0.0001, t(6)=11.16). (**d**) Dex anesthetized microglia have a greater number of maximum process intersections (n=4m/3f, Paired t-test, P=0.0286, t(6)=2.865). (**e**) Max projection of microglial processes over the hour imaging session to observe parenchyma surveillance. (**f**) Microglia in dex anesthetized animals survey a greater portion of the CNS parenchyma over an hour of imaging (n=4m/3f, Paired t-test, P=0.0204, t(6)=3.128). (**g**) Dex anesthesia decreases microglial process motility (n=4m/3f Paired t-test, P=0.0005, t(6)=6.867). (**h**) Dex anesthesia decreases microglial process instability (n=4m/3f Paired t-test, P=0.0008, t(6)=6.231). (**i**) Dex anesthesia increases microglial process stability (n=4m/3f Paired t-test, P<0.0001, t(6)=10.02). Scale bars 20μm. Graphs show mean ± s.e.m. Individual points represent individual animals.

### Anesthesia facilitates the microglial response to focal tissue injury

Given the significant impact of anesthesia on microglial surveillance and morphology, we next wanted to establish if these changes could affect targeted microglial responses to acute injury. To test this possibility, we generated focal laser ablation injuries in the cortex of the same mice while they were either awake or anesthetized, and quantified the microglial injury response of microglia in the immediate surrounding tissue over an hour (Fig.3a,b; Supplementary Videos 4 and 5). We found that anesthesia significantly increased the microglial response to focal tissue injury (Fig.3c-e; AUC; Paired t-test; p<0.05; Max response; Paired t-test; n.s.). The enhanced injury response and increased basal stability in anesthetized conditions suggest that wakefulness exerts a primarily inhibitory effect on microglial dynamics that is alleviated by sedation.

**Figure 3.**
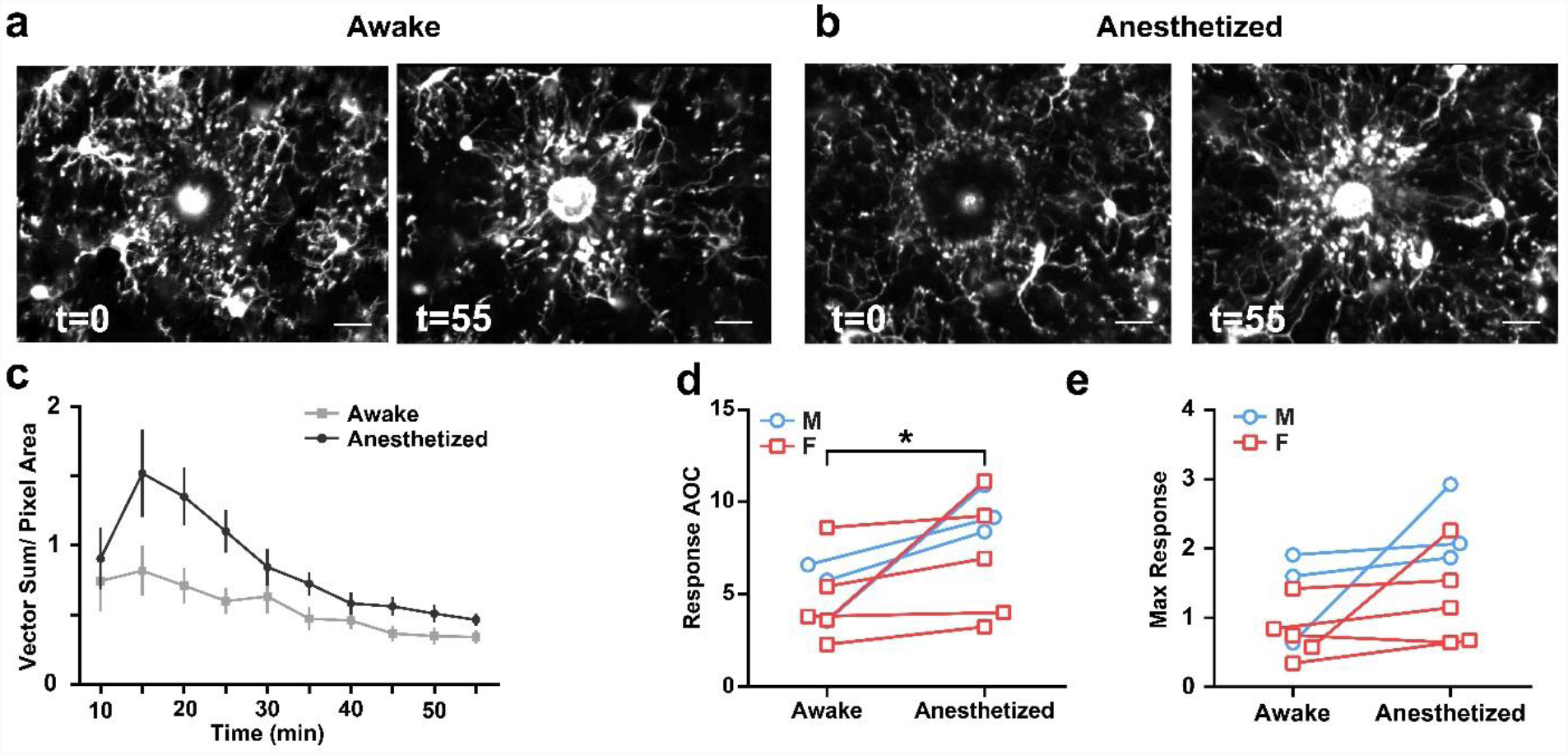
Anesthesia facilitates microglial process recruitment to focal tissue injury. (**a**) Microglia in an awake mouse surrounding a focal laser ablation injury at t=0 and t=55 minutes post-injury. (**b**) Microglia in an anesthetized mouse surrounding a focal laser ablation injury at t=0 and t=55 minutes post-injury. (**c**) Graph of microglial process recruitment from 10 minutes post-injury to 55 minutes post-injury. (**d**) Area under the curve of microglial process recruitment from 10-55min post-injury (n=3m/5f Paired t-test, P=0.0249, t(6)=2.844). (**e**) Microglia trend towards a greater magnitude of injury response in anesthetized conditions (n=3m/5f Paired t-test, P=0.0249, t(6)=2.844). Scale bars 20μm. Graphs show mean ± s.e.m. Individual points represent individual animals.

### β_2_ Adrenergic receptor signaling and noradrenergic tone alter microglial dynamics

To dissect the underlying mechanisms responsible for our observations, we considered signaling pathways in microglia that are likely to be dependent on wakeful states and that could affect microglial process stabilization. One prominent candidate is NE because DEX treatment decreases cortical NE levels, which could explain the microglial effects we observe following DEX administration. Furthermore, NE is a powerful mediator of wakefulness with decreased concentrations in the cortex during anesthesia^19,22^. NE exerts diverse effects on multiple cell types through a family of G-protein coupled receptors. Though it is possible that NE exerts its effects on microglia through indirect pathways involving other cell types, we focused on the direct effect on β2-ARs because microglia are known to express β2-ARs at higher levels than other CNS cell types in the healthy brain.^20^ In addition, previous work *in vitro* work has demonstrated that microglial β_2_-AR signaling can inhibit chemotaxis towards ATP^23^. To test whether β_2_-AR signaling alters microglial dynamics *in vivo*, we systemically applied the blood-brain barrier (BBB) permeant, β_2_-AR selective agonist, clenbuterol. To account for indirect clenbuterol effects through the periphery, mice were pre-dosed with the BBB impermeant, non-selective β-antagonist, nadolol at least 1 hour prior to clenbuterol dosing. Clenbuterol treatment caused a significant and sustained retraction of microglial processes, resulting in microglia that resemble those seen in awake mice (Fig.4a). Microglial pseudopodia retracted within the first 30min of imaging and remained retracted for the second half-hour of imaging (Fig.4b; Two-way ANOVA; p<0.0001). The retraction of pseudopodia was accompanied by a concomitant loss of *de novo* pseudopodia formation from 0-30min (Supplementary Fig.3). In addition, we also observed a persistent decrease in microglial motility as compared to saline-treated mice (Fig.4c; Two-way ANOVA; p<0.01). In order to further validate that the effects of β_2_-AR agonism on microglia were a direct effect on microglia and not due to peripheral activation of β_2_-ARs, we directly applied a selective β_2_-AR agonist, terbutaline, through an acute craniotomy and replicated our effects from the i.p. clenbuterol administration (Supplementary Fig.4, and Supplementary Video 6).

In order to further quantify the impact of clenbuterol on microglia we traced individual microglia and used Sholl analysis to assay the complexity of the arbor. We found that microglia treated with clenbuterol two hours before imaging had significantly reduced arbor complexity (Fig.4d-g; AUC; one-way ANOVA; p<0.0001; Maximum # of intersections; one-way ANOVA; p<0.0001). Clenbuterol-treated animals also had a significant decrease in microglial process coverage of the parenchyma over one hour (Fig.4h,i; one-way ANOVA; p<0.0001). We also wanted to assess if our observed effects of acute clenbuterol administration produced sustained changes in microglial dynamics beyond the initial hour post-administration. Examining microglial motility, we found it was significantly reduced in clenbuterol-treated animals (3 hours post-administration), although the effect was significant only in female mice (Fig. 4j; Two-way ANOVA; p<0.01).

**Figure 4.**
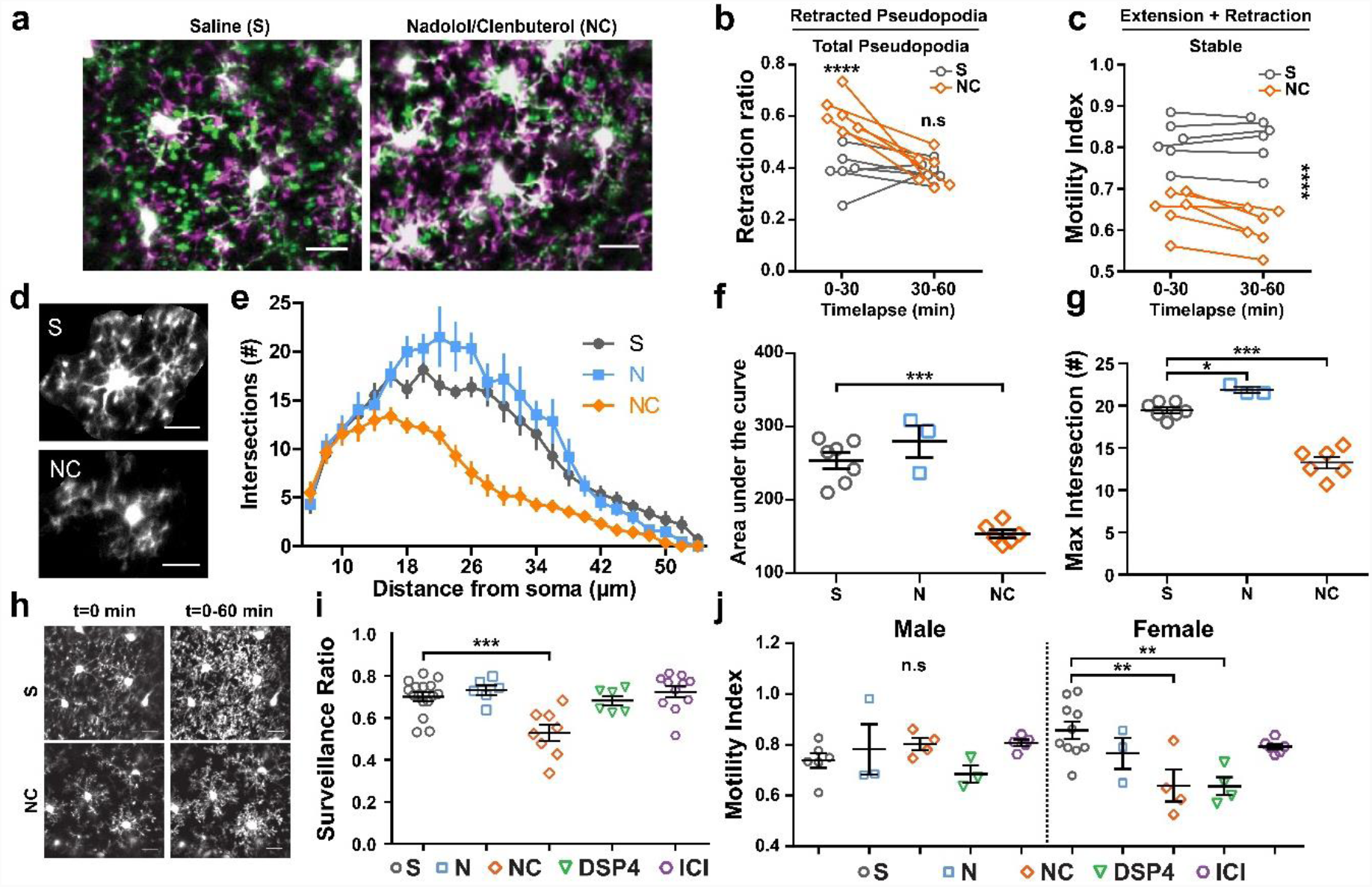
β_2_AR signaling profoundly reduces microglial dynamics in adolescent mice. (**a**) Overlay of t=0 (purple) and t=30 (green) of microglia from mice receiving either saline or clenbuterol at t=0 of imaging, note a robust degree of retraction in the clenbuterol treated mice.(**b**) Retraction ratio (retracted pseudopodia/total pseudopodia) Microglia treated with clenbuterol have a rapid transient increase in pseudopodia retraction from 0-30min (n=6 S, and n=6 NC, Two-way repeated measures ANOVA, P=0.0039, F(1,10)=13.91, Bonferroni post-tests for multiple comparisons, p<0.0001 s v. nc 0-30min). (**c**) Clenbuterol treated microglia have reduced process motility (n=3m/3f S, and n=3m/3f NC, Two-way repeated measures ANOVA, P=0.0037, F(1,10)=14.20, Bonferroni post-tests for multiple comparisons, p<0.0001 s v. nc 0-30min and 30-60min). (**d**) Examples of microglia from saline and clenbuterol treated animals. (**e**) Sholl profile of saline, nadolol, and nadolol/clenbuterol treated mice, note the robust reduction in the profile of clenbuterol treated mice. (**f**) Clenbuterol greatly reduces microglial arborization, this is reflected in a reduced area under the curve of the sholl profile (n=3-6 per group, One-way ANOVA, P<0.0001, F(2,15)=33.05, Dunnett’s multiple comparisons tests, S v. NC P<0.001). (**g**) The maximum number of intersections in the sholl profile is robustly reduces in clenbuterol treated microglia (n=3-6 per group, One-way ANOVA, P<0.0001, F(2,15)=61.13, Dunnett’s multiple comparisons tests, S v. N P<0.05, S v. NC P<0.001). (**h**) Microglial surveillance from 0-60min of motility imaging, note the lack of surveillance in the clenbuterol treated sample. (**i**) Microglia in clenbuterol treated mice survey a less of the parenchyma (n=5-17 per group, One-way ANOVA, P=0.0003, F(4,47)=6.737, Dunnett’s multiple comparisons tests, S v. NC P<0.001). (**j**) Microglial motility is reduced in NC-dosed and DSP4-dosed female mice (n=5-17 per group, One-way ANOVA, P=0.0108, F(4,39)=3.781, Dunnett’s multiple comparisons tests, S v. NC P<0.001 and S v. DSP4 P<0.001). Scale bars 20μm. Graphs show mean ± s.e.m. Individual points represent individual animals.

To further dissect the effects of β_2_-AR signaling on microglial dynamics, we assayed microglial motility in animals treated with different pharmacological manipulations of β_2_-AR or NE signaling. We asked whether reducing microglial β_2_-AR signaling affected microglial dynamics. We administered the β_2_-AR antagonist, ICI-118,551 (at time of imaging) to selectively block β_2_-AR signaling and depleted cortical NE using N-(2-chloroethyl)-N-ethyl-2-bromobenzylamine (DSP4, a neurotoxin selective for noradrenergic neurons). Neither treatment affected microglial surveillance (Fig.4h,i), although DSP4 reduced microglial motility in female mice (Fig.4j; Supplementary Movies 7-11; two-way ANOVA; p<0.05). The reductions in motility were associated with decreased instability in females and enhanced stability in both males and females (Supplementary Fig.5).

### Effects of modulating β_2_ -ARs in awake mice

Because β_2_-AR stimulation decreased microglial ramification and decreased motility in anesthetized animals, we next asked whether blocking or stimulating β_2_-AR signaling in awake mice affected microglial morphology and dynamics. Microglia imaged through cranial windows in awake mice were treated with either clenbuterol or ICI-118,551 to selectively stimulate or block β_2_-AR signaling respectively (Fig. 5a). Sholl analysis revealed that clenbuterol had no effect on microglial ramification in awake mice (Fig.5a-d; AUC; Paired t-test; n.s.; Maximum # of intersections; Paired t-test; n.s.). However, there was a significant increase in microglial ramification in ICI-118,551-treated animals as evidenced by an increased area under the Sholl curve and increase in max intersections (Fig.5a-b,e-f; AUC; Paired t-test; p<0.001; Max response; Paired t-test; p<0.0001). We found that clenbuterol significantly reduced microglial surveillance of the parenchyma, despite the lack of impact on microglial arborization (Fig.5g,h; Paired t-test; p<0.01). We also found that ICI-118,551-treated microglia showed enhanced surveillance of the parenchyma (Fig.5g,I; Paired t-test; p<0.05). In addition, we examined motility and found that clenbuterol significantly reduced microglial motility in awake mice (Fig.5j; Paired t-test; p<0.05), and that ICI-118,551 significantly increased microglial motility in awake mice (Fig.5k; Supplementary Video 12; Paired t-test; p<0.05;). Combined with our previous data in awake vs. anesthetized mice, these data suggest that during wakeful conditions, microglial ramification and process motility are inhibited by endogenous norepinephrine, and that relieving this inhibition via β_2_-AR antagonism induces an increase in process ramification and surveillance recapitulating the effects of anesthesia.

**Figure 5.**
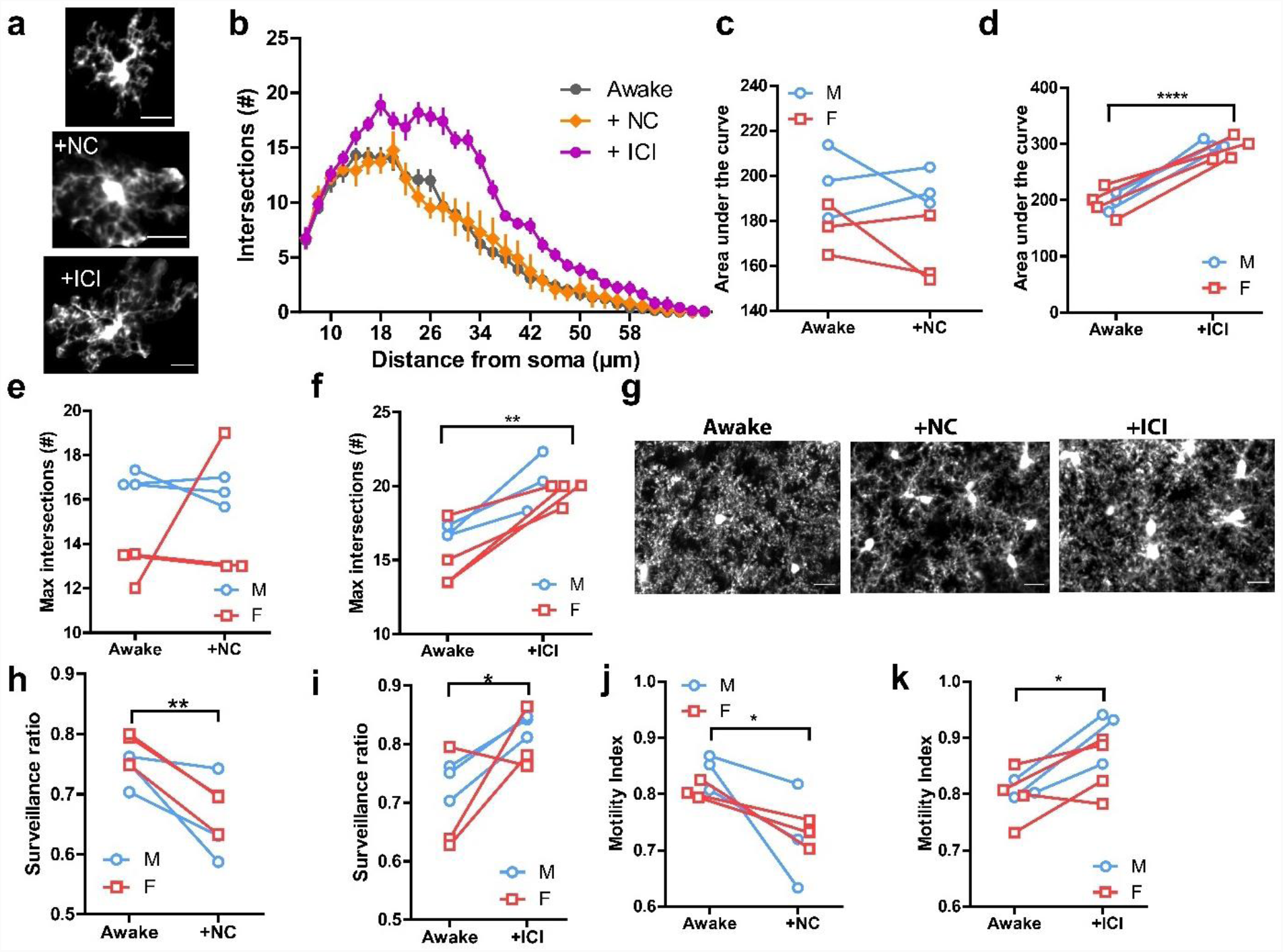
Inhibition of β_2_ ARs in awake mice recapitulates the effects of anesthesia by enhancing microglial arborization and surveillance. (**a**) Microglia from awake mice and NC or ICI-dosed awake mice. (**b**) Sholl profile of awake mice and NC or ICI-dosed awake mice, not the robust increase in arborization with ICI administration. (**c**) The area under the curve of awake mice given clenbuterol is not altered (n=3m/3f, Paired t-test, P=0.3642, t(5)=0.9977, n.s.) (**d**) ICI increases the area under the curve of the sholl profile, inhibition of β_2_ARs increases microglial arborization in awake mice (n=3m/4f, Paired t-test, P=0.0019, t(6)=5.244). (**e**) Maximum intersection # of the sholl profile is not altered with NC administration in awake mice (n=3m/3f, Paired t-test, P=0.5980, t(5)=0.5626, n.s.). (**f**) Maximum intersection # of the sholl profile is robustly increased by ICI inhibiting β_2_ARs (n=3m/4f, Paired t-test, P<0.0001, t(6)=14.89). (**g**) Microglial surveillance from 0-60min of motility imaging, note the decreased surveillance with clenbuterol treatment and enhanced surveillance with ICI treatment. (**h**) Microglia in clenbuterol-treated awake mice survey less of the parenchyma (n=3m/3f, Paired t-test, P=0.0045, t(5)=4.890) (**i**) Microglia in ICI-treated awake mice have enhanced surveillance of the parenchyma (n=3m/3f, Paired t-test, P=0.0290, t(5)=3.031) (**j**) In awake mice clenbuterol driven agonism of β_2_ARs reduces microglial motility (n=3m/3f, Paired t-test, P=0.0138, t(5)=3.711). (**k**) Microglial process motility is increased in ICI-dosed mice (n=3m/4f, Paired t-test, P<0.0156, t(6)=3.342). Scale bars 20μm. Graphs show mean ± s.e.m. Individual points represent individual animals.

### β_2_-AR signaling impacts microglial response to focal tissue injury

Previous work has shown that both NE and β-AR stimulation blocks microglial process chemotaxis *in vitro*^23^ suggesting that β-AR signaling may attenuate microglial responses to injury. To determine whether this is also true *in vivo*, we created a focal laser tissue injury in adolescent mice and observed reduced microglial process recruitment to the lesion core in clenbuterol treated mice (Fig.6a,b). We observed that peak response and overall microglial response to the injury was significantly reduced in clenbuterol-treated mice (Fig.6c-e; One-way ANOVA; p<0.0001). Furthermore, there was a trend towards and increased response in nadolol-or ICI-118,551-treated mice (Fig.6d; One-way ANOVA; p<0.0001). These data suggest that microglial β_2_-AR signaling attenuates not only baseline surveillance, but also the response to changes in CNS homeostasis during acute injury.

**Figure 6.**
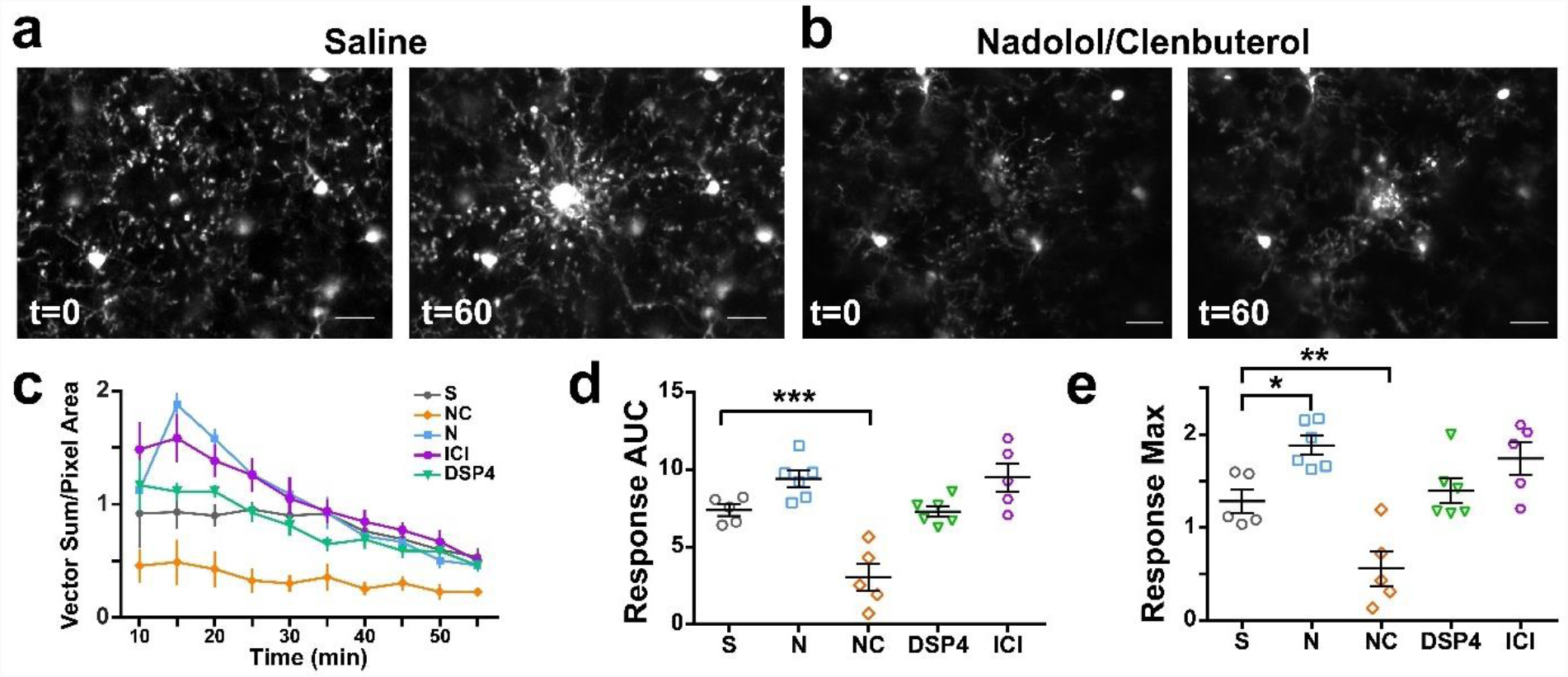
β2AR activation inhibits microglial process chemotaxis towards focal tissue injury. (**a**) Microglia in a saline-dosed mouse surrounding a focal laser ablation injury at t=0 and t=55 minutes post-injury. (**b**) Microglia in a clenbuterol-dosed mouse surrounding a focal laser ablation injury at t=0 and t=55 minutes post-injury. (**c**) Graph of microglial process recruitment from 10 minutes post-injury to 55 minutes post-injury (n=5-6 per group). (**d**) Area under the curve of microglial process recruitment from 10-55min post-injury, not the reduced response in clenbuterol-treated mice (n=5-6 per group One-way ANOVA, P<0.0001, F(4,26)=16.65, Dunnett’s multiple comparisons, S v. NC p<0.001). (**e**) Clenbuterol-treated microglia have a reduced maximum response to focal tissue injury (n=5-6 per group One-way ANOVA, P<0.0001, F(4,26)=12.69, Dunnett’s multiple comparisons, S v. NC p<0.01, S v. N p<0.05). Scale bars 20μm. Graphs show mean ± s.e.m. Individual points represent individual animals.

### Noradrenergic signaling and microglial β_2_-AR stimulation impair critical period ocular dominance plasticity

After observing the robust effects of noradrenergic and β_2_-AR signaling on microglia, we asked whether this signaling pathway mediates microglial roles in synaptic plasticity. Previously published work in our lab has shown that microglial processes interact with neural elements *in vivo*^*^4,7,10^*^ and that these interactions can be modulated by visual experience^4^. We have also shown that during the mouse adolescent visual critical period perturbation of microglial purinergic signaling through P2Y12 impairs ocular dominance plasticity (ODP)^10^. Given the antagonistic effect of β_2_-ARs on P2Y12 signaling^23^, we hypothesized that microglial β_2_-AR signaling might impair ODP during the peak of the mouse visual critical period (∼P25-P32^24,25^), when our experiments have demonstrated altered microglial phenotypes (Fig. 4). Selectively activating β_2_-ARs chronically with clenbuterol effectively blocked ODP, indicated by the lack of ocular dominance shift of neuronal responses after four days of monocular deprivation (Fig.7b;Two-way ANOVA; p<0.001). As before, nadolol was administered with clenbuterol to block peripheral β_2_-AR stimulation. Both saline-treated and nadolol-only treated animals showed normal ocular dominance shifts indicating intact ODP (Fig.7a,b). In addition, cortical NE depletion with DSP4 treatment impaired ODP confirming previous work demonstrating that NE has important roles in ODP (Fig.7b)^26^. However, this effect appeared to be independent of β_2_-AR signaling as chronic blockade of β_2_-ARs with ICI-118,551 did not impact ODP (Supplementary Fig.6). Thus, sustained stimulation of β_2_-AR signaling interferes with the plastic processes that mediate ODP, but NE signaling through other signaling pathways are necessary for ODP^27^.

### Microglial β_2_-AR activation impairs ocular dominance plasticity

Despite the known roles of microglia in ocular dominance plasticity and high microglial β_2_-AR expression, it is possible that pharmacological β_2_-AR stimulation impairs ODP through non-microglial pathways. To determine whether our results are specifically due to β_2_-AR signaling in microglia, we crossed the CX3CR1Cre-ERT^9^ mouse line to β_2_-AR floxed mice^28^ to selectively ablate β_2_-ARs in microglia. Mice were dosed with tamoxifen from ∼p2-4^9^ to ensure that β_2_-ARs were selectively ablated from microglia in adolescent mice. Microglia were isolated from tamoxifen-treated animals and successful gene excision of β_2_-ARs confirmed by PCR (Supplementary Fig.7). To test whether the effects of β_2_-AR stimulation on microglial dynamics were dependent on microglial β_2_-AR expression, we labeled microglia by crossing these mice with the tdTomato-labeling, Ai9 line. Clenbuterol no longer induced a retraction of the microglial arbor in tamoxifen-treated Cx3cr1Cre-ERT-Het/β_2_-ARflx/Ai9 mice, while the response was intact in control, tamoxifen-treated Cx3cr1Cre-ERT-Het/Ai9 mice (Fig. 7c,d; Two-way ANOVA; p<.0001), further confirming loss of these receptors from microglia. We then repeated ODP experiments with tamoxifen-treated Cx3cr1Cre-ERT-Het controls and Cx3cr1Cre-ERT-Het/β_2_-ARflx (Cx3cr1Cre/β_2_-ARflx) mice (Fig.7e-h). Control mice showed the same effects of pharmacological manipulation as C57BL/6J mice (Fig.7f; Two-way ANOVA; p<0.05). Cx3cr1Cre/β_2_-ARflx mice lacking microglial β_2_-ARs showed normal ODP, suggesting that microglial β_2_-ARs are not necessary for ODP. We found that ablating β_2_-ARs in microglia, however, rescued ODP during chronic clenbuterol administration (Fig.7g,h; Two-way ANOVA; n.s. interaction), suggesting that aberrant activity of microglial β_2_-ARs interferes with normal plasticity. DSP4 treatment still blocked ODP in the Cx3cr1Cre/β_2_-ARflx mice (Fig.7h; Two-way ANOVA; Bonferroni post-hoc; p<0.05), showing that the effects of depleting NE are mediated by other cell types. Thus, we confirmed that chronic stimulation of microglial-specific β_2_-ARs is capable of blocking ODP while cortical NE depletion likely blocks ODP through non-microglial mechanisms.

**Figure 7.**
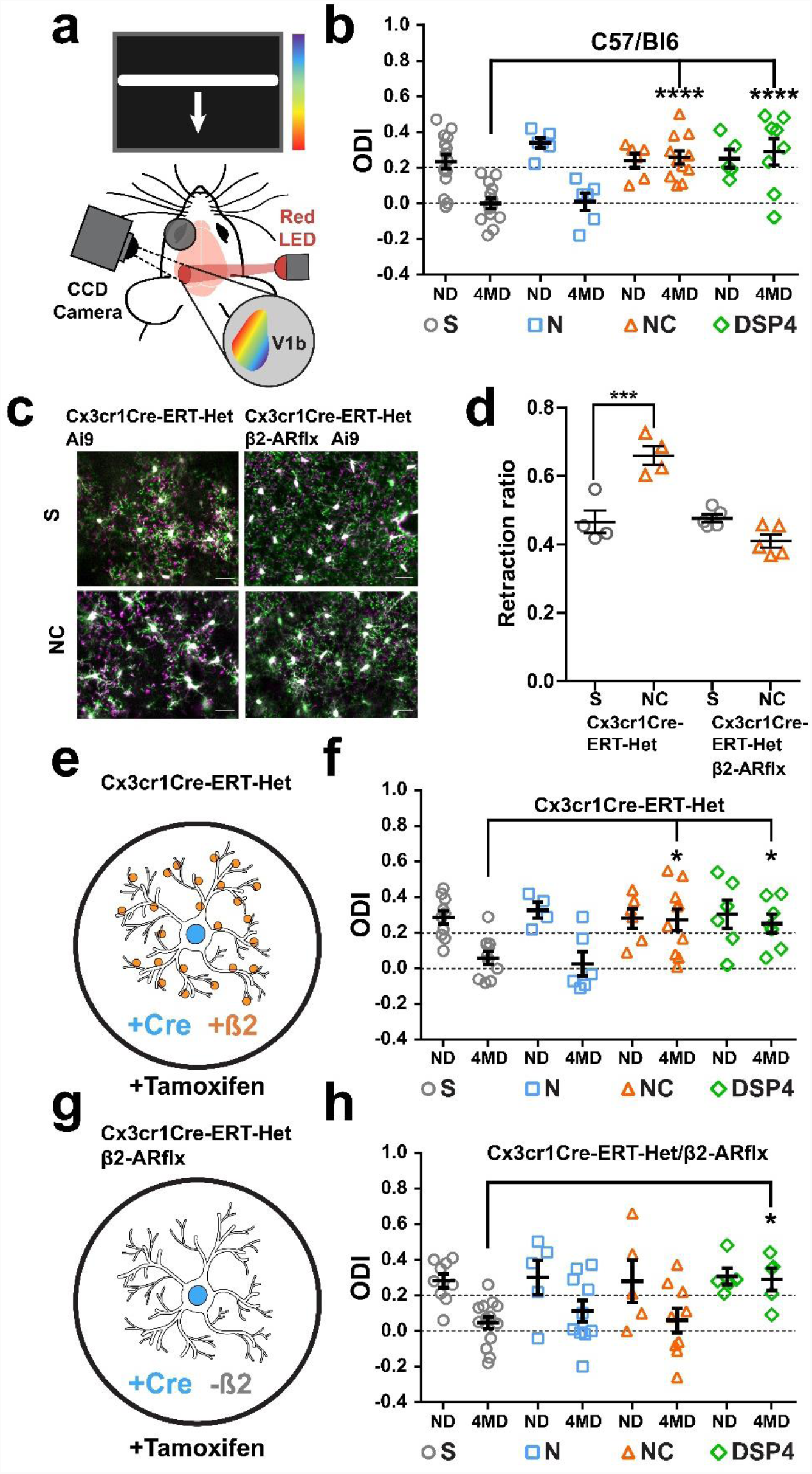
Chronic microglial β_2_ AR activation impairs adolescent ocular dominance plasticity. (**a**)Schematic of the intrinsic optical signal-imaging technique used in our experiments (**b**) Quantification of ocular dominance indices demonstrates robust shifts in saline and nadolol treated groups, but no shifts in Clenbuterol and DSP4 treated groups (n=5-15 per group, two-way ANOVA, P=0.0005, F(3,63)=6.809, Bonferroni multiple comparisons, S 4MD v. NC 4MD p<0.0001, S 4MD v. DSP4 4MD). (**c**) Overlay of Cx3cr1-CRE/Ai9 and Cx3cr1-CRE/B2/Ai9 microglia at t=0 and t=30 in saline or clenbuterol-treated mice, note the lack of process retraction with the ablation of microglial β_2_ARs. (**d**) Selective ablation of microglial β_2_ARs rescues microglial pseudopodia retraction in clenbuterol-dosed mice (n=4-5 per group, Two-way ANOVA, P<.0001, F(1,14)=32.11, CreAi9S v. CreAi9NC p<0.001).(**e**) Schematic of Cx3cr1-CRE microglia representing intact microglial β_2_ARs. (**f**) Quantification of ocular dominance indices in Cx3cr1-CRE mice demonstrates robust shifts in saline and nadolol treated groups, but no shifts in Clenbuterol and DSP4 treated groups (n=4-10 per group, Two-way ANOVA, P<0.0346, F(3,52)=3.099, Bonferroni multiple comparisons, S 4MD, v. NC 4MD p<0.05, S 4MD v. DSP4 4MD p<0.05). (**g**) Schematic of Cx3cr1-CRE/B2 microglia representing ablated microglial β_2_ARs. (**h**) Quantification of ocular dominance indices in Cx3cr1-CRE/B2 mice, ablation of microglial β_2_ARs rescues plasticity in clenbuterol-treated mice, but not DSP4-treated mice (n=5-14 per group, Two-way ANOVA, P<0.0008, F(1,54)=12.62, Bonferroni multiple comparisons, S 4MD, v. DSP4 4MD p<0.05).

## Discussion

We report that the microglial β_2_-AR is a key receptor in regulating basal microglial surveillance and injury response in the visual cortex of awake mice, and that chronic pharmacological stimulation of this receptor during the mouse adolescent visual critical period impairs cortical experience-dependent plasticity. In wakeful conditions, microglial process arbors are small and highly motile, dominated mostly by filopodial protrusions. Anesthesia rapidly increases microglial arborization and pseudopodia stabilization allowing microglial processes to survey a larger area of cortex and respond more rapidly to a focal injury. Administration of the β_2_-AR agonist clenbuterol in anesthetized mice results in microglial characteristics that resemble the awake state, while the β_2_-AR antagonist ICI-118,551 generates microglial phenotypes that resemble anesthetized conditions in awake mice, suggesting that signaling through this microglial receptor regulates microglia function during wakefulness. Our findings indicate that there are robust differences in microglial basal dynamics between awake and anesthetic states and that these changes are driven by noradrenergic β_2_-AR signaling in microglia (Fig.8).

### Endogenous NE is a potent modulator of microglial function

NE is a potent neuromodulator that regulates brain function and elicits transitions between internal states such as wakefulness, arousal, and stress. NE has many targets throughout the nervous system and can coordinate synchronized changes in multiple cell types through both tonic signaling during wakefulness and phasic firing to salient or noxious stimuli.^19^ Astrocytic calcium signaling is robustly regulated by NE,^15,29^ which can impact potassium buffering, neurovascular coupling, neural metabolism, and neurotransmitter recycling.^30^ Astrocytic function is also highly dependent on brain state with different roles during sleep, when NE is low, and wakefulness or stress, when NE is high^22^. Here we provide evidence that microglia are another target for NE signaling. Our work suggests that microglia, like astrocytes, may differ robustly between the awake and anesthetized or “sleep-like” state. While we could not quantify microglial dynamics in sleeping animals, due to the fact that mice have very short sleep cycles^22,31^, our experiments with dexmedetomidine suggest that microglia may also show enhanced surveillance and injury responses during sleep. Dexmedetomidine is thought to hyperpolarize LC neurons and inhibit NE release throughout the brain^21^, leading to brain activity that resembles more prolonged slow-wave sleep in humans^31^. Thus, our work indicates that microglia may have different functions in the awake vs. sleeping brain. This idea merits further investigation, as many diseases are linked to changes in sleeping patterns, and the quality of sleep can in turn greatly impact disease progression^32^.

Noradrenergic inhibition of microglial dynamics impairs synaptic plasticity, suggesting that brain state, through noradrenergic signaling, modulates microglia-synapse interactions^4^. Simulating a chronic awake state through pharmacological targeting of microglial β2-ARs prevents ODP, further suggesting that microglia require a release from noradrenergic signaling in order to fulfill essential roles in neural plasticity. This implies that in physiological conditions NE may be exerting an important inhibitory function on microglia, perhaps preventing inappropriate or excessive microglia-neuron interactions. In the awake behaving state tonic NE release decreases neural variability in firing rates and is thought to enhance the signal to noise ratio in the visual cortex^33^. NE bulk release from varicosities could enhance neural tuning while simultaneously inhibiting microglial-neuron interactions through selective activation of β_2_-ARs. Previous work has demonstrated that sleep is necessary for plasticity, as well as specifically for ODP^34^, and while a multitude of signals are regulated by sleep, microglial β_2_-AR signaling may be an additional target. It is also important to note that microglial P2Y12 is necessary for ODP and both genetic and pharmacological inhibition results in the loss of plasticity^10^. Thus, it is likely that interactions between β_2_-AR and P2Y12 signaling in microglia regulate microglial roles at synapses. Effects of NE on microglia and plasticity may also be mediated by NE signaling in other cell types. In our study, depletion of NE with the neurotoxin DSP4 blocked ODP and affected microglial motility, however ablation of microglial β_2_-ARs failed to rescue plasticity in DSP4 treated mice. This suggests that loss of NE in this model signals through a different receptor or cell-type to alter microglial dynamics and plasticity. Interestingly, astrocytes have robust expression levels of α _1_, α _2_-, and β_1_, ARs which contribute to astrocytic metabolic coupling and K+ clearing during neural activity^19^. The diverse effects of NE on astrocytic function could make astrocytes an intermediary by which depletion of NE may be altering both microglial function and neural plasticity. It is also important to note that selective β_2_-AR signaling does not account for all changes in microglial dynamics. Wakefulness and β_2_-AR stimulation had opposite effects on microglial motility suggesting that motility *per se* is regulated through a different signaling pathway possible initiated by NE signaling in other cell types. The regulation of glymphatic flow by wakefulness may be an additional mechanism by which NE signaling in other cell types may influence microglial dynamics, in this case by altering extracellular space^16^.

### Intracellular signaling effectors of β_2_-AR in microglia

β_2_-AR and P2Y12 signaling have opposite effects on microglial morphology, chemotaxis^35^ and ODP. Both receptors are G-protein coupled. While P2Y12 is coupled to the G_i_ subunit, the β_2_ AR is coupled to G_S_, suggesting that these may act in a push-pull system^*^23^*^ (Fig. 8). The intracellular pathways that mediate G protein signaling in microglia are still unclear largely because reduced preparations do not necessarily capture physiological microglial behavior. E*x vivo*, β_2_-AR stimulation by NE and β_2_-AR agonists blocks P2Y12 process chemotaxis toward ATP by downstream protein kinase A (PKA) dependent phosphorylation of phosphoinositide 3-kinase gamma (PI3Kγ)^23,36^, suggesting this pathway may be important. Push-pull signaling through adenylyl cyclase may also play a role. Recent studies in the acute brain slice have shown that baseline microglial motility relies on the maintenance of the microglial resting membrane potential by the two-pore domain channel THIK-1^37,38^. Genetic or pharmacological perturbations of THIK-1 function reduce microglial ramification and surveillance, similarly to β_2_-AR stimulation, making it a likely target. While P2Y12 signaling does not alter basal surveillance in microglia, it does gate THIK-1 function causing hyperpolarization in response to ATP. β_2_-ARs could similarly interface with THIK-1 or Kir K^+^ channels to alter the microglial resting membrane potential thereby inhibiting surveillance. This is the case in neurons of the medial prefrontal cortex, where clenbuterol hyperpolarizes fast-spiking GABAergic interneurons^39^ due to β_2_-AR driven Gαs inhibition of inward Kir K^+^ rectifying channels. β_2_-ARs could also regulate microglial motility through recruitment and signaling by β-arrestin1 and 2 which are both expressed in mouse microglia^40^ and can have diverse downstream targets including cytoskeletal proteins^41^. Recent work in zebrafish showed a critical role of β-arrestin1 in maintaining normal microglial morphology, surveillance, and phagocytosis^42^. Long-term activation of β_2_-ARs can lead to heterologous desensitization and possible endocytosis of other GPCRs such as P2Y12^41^ through β-arrestins. Activation of β-arrestin2 in microglia initiates longer term anti-inflammatory signaling through blockade of Mitogen-activated protein kinases (MAPKs) rescuing neuronal death in a Parkinson’s disease model^43^. P2Y12 signaling can also drive IL-1β and inflammasome recruitment in response to LPS triggered inflammation^37^, suggesting that cytokine effectors may critically alter microglial and inflammatory milieu after activation of GPCRs. It is important to note that many forms of plasticity^44^, including ODP^45^, are thought to be mediated by cytokines, such as IL-1 β and TNFα, which are also known to promote sleep^46^. Our data suggests that microglial β_2_-AR signaling may be regulated by sleep-wake cycles. Microglial β_2_-ARs may in turn also alter sleep through regulation of cytokine production driven by downstream nuclear factor ϰβ signaling^47^. From previous work and our findings, a picture emerges of a delicate interplay between P2Y12 and β_2_-AR signaling which mediate the responsivity of microglia to acute perturbations in the CNS milieu.

**Figure 8.**
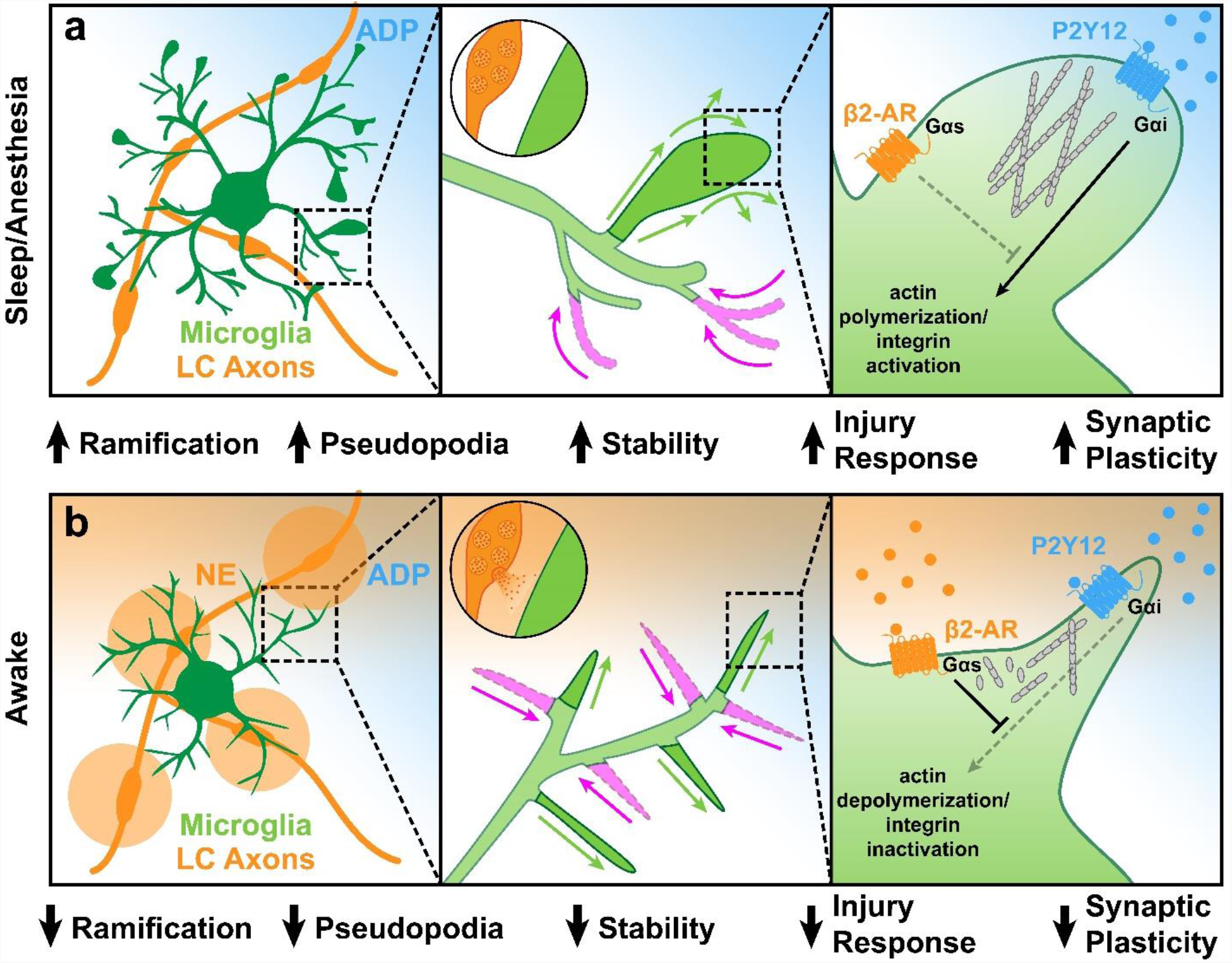
Summary model of results. (**a**) Microglial dynamics during sleep/ anesthesia. Left panel: during sleep-like states/ anesthesia microglia (green) extend secondary and tertiary processes to adopt a more ramified state when cortical norepinephrine release from locus coeruleus axons (orange) are low. Larger pseudopodia extend and remain stable. Middle panel: microglial motility decreases as processes become more stable (green arrows) and filopodia retract (magenta). Pseudopodia formation increases towards chemokinetic factors such as ADP, increasing microglial responses to injury and facilitating microglial roles in synaptic plasticity. Right panel: Gαi-coupled signaling (such as from P2Y12) activates integrins and facilitates actin polymerization within extending pseudopodia. (**b**) Microglial dynamics during wakeful states. Left panel: during wakeful states, cortical norepinephrine (NE) release from locus coeruleus axons (LC, orange) cause microglial secondary and tertiary processes (green) to retract reducing microglial process ramification and reduces the formation of pseudopodia. Middle panel: microglial motility increases as processes are less stable and filopodia quickly extend (green) and retract (magenta). Right panel: Gαi-coupled signaling is attenuated by norepinephrine via Gαs-coupled β2-ARs resulting in integrin deactivation and actin depolymerization. Subsequently, microglial responses to injury and roles in synaptic plasticity are inhibited.

### Implications for understanding microglial signaling in disease

The diversity of intracellular signaling pathways down-stream of microglial β_2_-ARs make this receptor a fascinating target for the gating of microglial physiology and responses. Our results demonstrate profound effects of selective β_2_-AR stimulation and wakefulness on microglial behavior. These mechanisms may also have a large impact on disease processes. It has been well documented that sleep disturbances are a hallmark and some of the earliest indicators of the onset of neurodegenerative diseases such as Parkinson’s disease (PD), Alzheimer’s disease (AD), and Huntington disease (HD)^48^. In the case of AD, sleep deprivation and sleep loss lead to elevated levels of pathological tau and β-amyloid (Aβ), thereby worsening neural health and synapse loss. Interestingly, prolonged wakefulness and increased noradrenergic signaling prevent adequate astrocyte-driven glymphatic clearance of Aβ ^16^. Sleep loss in AD could initiate a range of deleterious processes by increasing noradrenergic signaling: a failure of astrocytic clearance mechanisms permitting accumulation of toxic proteins such as Aβ with an accompanying failure of microglial surveillance and recruitment to sites of injury and degeneration due to enhanced β_2_-AR activity. On the other hand, NE has been shown to have beneficial roles on microglia in AD, where it enhances Aβ plaque clearance by microglia while simultaneously dampening the production of proinflammatory cytokines^49^. This is in line with other disease models. For instance, in a model of PD, β_2_-AR activation enhances microglial anti-inflammatory signaling rescuing neural cell death^43^. Similarly, in kainate-induced seizures, β_2_-AR activation promotes an anti-inflammatory phenotype in microglia decreasing damage and neural apoptosis^50^. While we found that acute noradrenergic signaling inhibits microglial recruitment to acute tissue damage, NE may be helpful in mediating microglial responses to chronic degenerative disease states. Further research is needed to characterize the conditions where β_2_-AR activation and suppression of microglial responses by promoting an anti-inflammatory phenotype are either beneficial or deleterious for disease resolution.

Altogether, our data show that microglial β_2_-ARs are potent regulators of microglial dynamics and visual system experience-dependent plasticity. While future work is needed to address how microglia change during physiological sleep processes, our study provides promising evidence that microglia have critically different roles in the sleeping and awake brain.

## Methods

### Animals

Experimental protocols were carried out in strict accordance with the University of Rochester and Massachusetts Institute of Technology Committees on Animal Resources and conformed to the National Institute of Health guidelines. Experiments were conducted on mice with a C57/Bl6 background between 3-8 weeks of age (P21-P80). CX3CR1-GFP^1^(JAX: 005582) heterozygous mice were used to visualize microglia for *in vivo* two-photon imaging. CX3CR1-Cre^ERT2^ (JAX: 021160) mice were bred to Adrbeta2-flox^3^(Karsenty Lab courtesy of the Rosen Lab) mice to generate mice which had β_2_ARs selectively knocked out of microglia. CX3CR1-Cre^ERT^ mice were also bred to Ai9^4^ (tdTomato JAX: 007909) mice for visualization of microglia in 2-photon experiments. Mice were housed with a standard 12:12 light/dark cycle and fed standard chow *ad libitum*. For experiments regarding ocular dominance plasticity, mice were used in the visual critical period (P25-35). Sex distribution is provided in Supplementary Table 1.

### Stress Exposure

For stress exposure experiments, male and female mice were randomly assigned to control or stressed cohorts. Stressed mice were briefly anesthetized using 3% isoflurane and restrained in 50ml conical tubes drilled with air holes in a brightly lit fume hood for 2 hours. This stress period was repeated for three consecutive days at approximately the same time each day. Following the third day, mice were either imaged immediately following or 4 hours after the final stressor. Control mice were handled on each day but returned to their home cages.

### Circadian Rhythm Measurements

For circadian rhythm experiments, mice were kept at the standard 12:12 light/dark cycle with lights on at 6:00 and lights off at 18:00. For fixed-tissue analysis, brains were collected at 6:00, 12:00, 18:00, and 24:00. For *in vivo* experiments, animals were imaged at 12:00 and 24:00. For collection times during the dark cycle, cages were covered with a black sheet to minimize light exposure before experiments.

### Flow isolation of microglia and confirmation of β_2_AR excision

We utilized our previously published experimental design^5^ with a small modification to accommodate for the YFP expression in the CX3CR1-Cre^ERT^ line. CX3CR1-Cre^ERT^/ Adrbeta2-flox mice were either dosed with 50μg tamoxifen from P2-4 or undosed. At ∼P28 mice were euthanized by an i.p. sodium pentobarbital overdose (Euthasol, Virbac) and transcardially perfused with ice cold 0.15 M PBS. Each brain was removed and the whole cortex was dissected from the rest of the brain in ice cold, degassed FACS buffer (0.5% BSA, Sigma A2153 in 1x PBS Invitrogen 20012-027 pH 7.2). The tissue was kept on ice throughout all the procedures. Cortices were homogenized with a Dounce homogenizer and then passed through a 70μm filter and centrifuged (210xg, 7min, 4 °C). The supernatant was aspirated from the pellets and then the pellets resuspended and prepared for magnetic sorting with Myelin Removal Beads II (Miltenyi 130-0960733). The resuspended labelled tissue was passed through a 70μm filter and then the FACS buffer primed Magnetic columns (Miltenyi 130-096-733). After passage through the magnetic columns the samples were centrifuged, the supernatant removed, and resuspended in FACS buffer with Fc block (Biolegend 101320) for 15min at 4 °C.

Samples were incubated with CD11b-AlexaFluor BV786 (BD Biosciences 740861) and CD45-APC (BD Biosciences 561018) for 30 min in the dark at 4°C. Each sort included the following compensation controls: BV720 bead control, APC bead control (eBiosciences 01-111-42), unstained live cells, and propidium iodide (PI) labelled Triton X-100 killed cells. An additional fully-labelled sample was prepared with spare tissue to check the voltage settings prior to running experimental samples. All samples were run on an 18-color FACS Aria II flow cytometer. For the sort, all samples and controls were resuspended in 300μl FACS buffer. PI was added to the samples just prior to sorting. An example of a microglial sort can be found in Supplementary figure 8.

Cells were collected in PBS for DNA isolation with a Qiagen DNAeasy DNA isolation kit. A total of 3 tamoxifen treated and 3 tamoxifen untreated animals were included in our confirmation experiments. Isolated DNA was run through 2 PCR protocols, 1^st^ the PCR for confirmation of floxed allele presence (550BP product: Fw: ccaaagttgttgcacgtcac, Rv: gcacacgccaaggagattat), and 2^nd^ excision confirmation(∼800BP: Fw: ccaaagttgttgcacgtcac, Rv: aagaaagaggaggggctgag)^3^. We confirmed that in tamoxifen treated mice the 550BP floxed allele PCR product was no longer present (Supplementary figure 8), and the excision product ∼800BP was present. In our untreated mice we still had the 550BP floxed allele, however we did also see the ∼800BP excision product in the second PCR. This suggests that there is a certain degree of leakiness in our Cre expression. However, this leakiness did not affect our experimental outcomes as our controls for all CX3CR1-Cre^ERT^/ Adrbeta2-flox experiments were CX3CR1-Cre^ERT^-HET mice dosed with tamoxifen.

### Pharmacological Agents

Fentanyl cocktail^6-9^: fentanyl (0.05mg kg^−1^, i.p.), midazolam (5.0 mgkg^−1^, i.p.), and dexmedetomidine (0.5 mgkg^−1^) Pre-mixed and given i.p. Used for anesthetized 2-photon imaging sessions and for thin-skull and cranial window procedures. Dexmedetomidine^10^: 0.4 mg kg^−1^, i.p. for 2-photon imaging.

Clenbuterol^11^ (Sigma, 21898-19-1): 1-5 mg kg^−1^, i.p. for 2-photon imaging and 5 mg kg^−1^, i.p. once every 12hrs for 4 days for iOS. Nadolol^12^(Sigma, 42200-33-9): 10 mg kg^−1^, i.p. for 2-photon imaging and once every 12hrs for 4 days for iOS. ICI 118,551^13^ (Sigma, 72795-01-8): 10 mg kg^−1^, i.p. for 2-photon imaging and 8 mg kg^−1^ day^−1^ by mini-osmotic pump for 4 days (Alzet, 1007D) for iOS. N--N-ethyl-2-bromobenzylamine (DSP4)^14^ (Sigma, 40616-75-9) : 50 mg kg^−1^, i.p. twice at 48hr intervals a minimum of 48 hours prior to either 2-photon imaging or to the deprivation period of iOS. Tamoxifen (Sigma, 10540-29-1): 50μg by intragastric gavage once daily from ∼post-natal day 2-4 in experiments utilizing CX3CR1-Cre^ERT^ mice^2^.

### Monocular deprivation

Animals were randomly assigned to ND or 4MD cohorts starting from P27+2. MD animals were anesthetized (isoflurane, 5% induction, 3% maintenance). The right eyelids were trimmed and anti-biotic eye ointment was applied to the eye and trimmed margins. The eye was then sutured closed. Mice were given monitored and given carprofen (5 mg kg^−1^) every 24 hours for analgesia as needed. The eye was not reopened until the day of imaging and any animals with a compromised eye or sutures were excluded from the rest of the experiment.

### iOS Imaging

These experiments utilized C57/Bl6, CX3CR1-Cre^ERT^-HET, and CX3CR1-Cre^ERT^-HET/ Adrbeta2-flox. Following 4 days of ND or MD, animals were re-anaesthetized with isoflurane and chloroproxithene (2 mg kg^−1^), and sutured eyes reopened. The skull over contralateral visual cortex was exposed, cleared, covered with agarose (0.25%) and sealed with a coverslip. Animal anesthesia was maintained with isoflurane (0.75%) throughout imaging. A custom-made iOS imaging set-up was used to record activity in the visual cortex during presentation of a visual stimulus (DALSA 2M30 CCD). The cortex was illuminated with 550-nm light to identify vasculature and 700-nm light for iOS collection. Images of the left visual cortex were collected continuously, while either the ipsilateral or contralateral eye was stimulated by white horizontal square-wave bar gratings on a black background moving upwards (90°) or downwards (270°) at a frequency of 8°/s for 6min (30 cm from eyes). Visually evoked responses were collected for each eye individually. The normalized amplitude of the fast Fourier transform component of the intrinsic signal was averaged for each eye from responses to both stimulus directions and compared between eyes offline using MATLAB to determine ocular dominance. An ODI was computed using the following equation: ODI=(average contralateral response-average ipsilateral response) / (average contralateral response / average ipsilateral response).

### Cranial Window Surgery

Animals were anesthetized using the fentanyl cocktail during the cranial window implantation surgical procedure. Lubricant ointment was used to keep the eyes moist and protected. Body temperature was maintained at 37°C during the surgery. Asceptic technique was adhered to during all surgical procedures: all tools were autoclaved for steam sterilization, and tools were sterilized in a bead sterilizer between surgeries (up to 3 uses). Mice were mounted in a stereotaxic frame and head-fixed for surgical procedures. The skull was exposed through a scalp incision and all connective tissues were cleared off the skull. A 3mm biopsy punch (Integra) was then used to create a circular score on the skull over V1. A 0.5 mm drill bit (FST) was used to then drill through the skull for the craniotomy, tracing the 3mm score. A 5mm coverslip attached to a 3mm coverslip (Warner Instruments) by UV glue (Norland Optical Adhesive, Norland Inc) was then slowly lowered down into the craniotomy (3mm side down). The coverslip was carefully secured with Loctite 404 glue (Henkel Corp). A custom headplate produced by emachine shop (www.emachineshop.com) (designs courtesy of the Mriganka Sur Lab, MIT) was then secured onto the skull utilizing C&B Metabond dental cement (Parkell Inc). The cement was then used to cover the rest of skull and seal the incision site. Animals were administered slow-release buprenex by URMC veterinary staff (s.q. mg kg^−1^ every 72 hours) and monitored for 72 hours post-op.

### Two-Photon Microscopy

A custom two-photon laser-scanning microscope was used for *in vivo* imaging (Ti:Sapphire, Mai-Tai, Spectra Physics; modified Fluoview confocal scan head, 20x lens, 0.95 numerical aperture, Olympus). Excitation for fluorescent imaging was achieved with 100-fs laser pulses (80MHz) at 920nm for GFP and 1020nm for tdTomato with a power of ∼40mW measured at the sample. For motility experiments in CX3CR1-GFP mice a 580/180 (GFP) filter was used. For experiments in CX3CR1-Cre+/Ai9 mice a 565 a 578/105 filter was used. During anesthetized cranial window imaging sessions mice were anesthetized with our fentanyl cocktail. Prior to awake imaging sessions mice were trained for three consecutive days in head-restraint on a running wheel progressing from 30 minutes to 1 hour of head-restraint over the three days. During awake imaging sessions mice were head-restrained and kept on a foam running wheel for the duration of the session, the restraint apparatus and wheel were the same as were used in the initial training sessions. During thin-skull ^6,7,9^ imaging sessions mice were anesthetized with the fentanyl cocktail prior to the skull-thinning surgery and for the duration of the imaging session. During imaging sessions and during post imaging recovery mice were kept at 37°C until they were alert. Imaging was conducted at 3-5 digital zoom and 1um z-step, with time-lapse imaging at 5-minute intervals over 1 hour. Image analysis was done offline using ImageJ and MATLAB with custom algorithms as described in Sipe et al. 2016 and available upon request.

#### Microglial Morphology

For *in vivo* morphological analysis 2-3 microglia were selected per animal from a single time-point in the imaging session. For consistency the same time-point was selected across all imaging sessions when analyzing morphology. For each microglia selected an individual z-projection was created in ImageJ, which encompassed the entire microglial arbor. All the microglial processes were manually traced and then the tracing of the processes was subjected to sholl analysis. For each animal an average of the 2-3 microglia selected was found and that average represented the value for that animal. The area under the curve of the Sholl profile and then maximum intersection number were found and analyzed as indices of microglial arbor complexity.

#### Microglial motility and surveillance

Microglial motility analysis was performed in ImageJ and MATLAB as previously described ^6,8^. Z-stacks were collected in V1 every 5 minutes for 1 hour, producing 12 time points. All z-stacks were between 60-120µm and for analysis uniform z-projections were made (30-40µm thick). Lateral motion artifact was corrected (Stackreg plugin, ImageJ). A custom MATLAB algorithm ^6,8^ was used to compare pixels across individual time points and across consecutive time points to generate a motility index (defined as the sum of all changed pixels by the unchanged pixels). Additional indices were generated by the program including stability and instability (Fig1). For each index, all microglia in the Z-projection of the imaging session were analyzed to generate the value per animal for the 1-hour session. The thresholded time points were also used to calculate the area monitored (surveillance measurement) by microglia during the 1-hour imaging session. This was done by collapsing the 12 time points through the z-projection function in ImageJ and calculating the total number of pixels representing microglia normalized to the number of pixels in the first time point.

#### Microglial pseudopodia measurements

For pseudopodia analysis t=0, t=30, and t=60 minutes from the motility imaging sessions were used to create 0-30-and 30-60-minute overlays. The number of pseudopodia were then counted using the multipoint tool in imageJ to track marked pseudopodia. Pseudopodia were defined as bulbous end-tips of microglial processes. Retraction and extension ratios were calculated based on the number of retracted or extended pseudopodia divided by the total pseudopodia counted. For imaging of Ai9 animals, only 0 and 30 minute timepoints were taken as the tdTomato fluorophore suffered from pronounced photobleaching with repeated imaging. This precluded the use of Ai9 for motility measurements.

#### Terbutaline Motility

For direct application of terbutaline to the brain parenchyma, small craniotomies were performed on anesthetized animals and imaged directly without a coverslip. Baseline imaging was collected every 2 minutes for 30 minutes in a sterile saline objective immersion. Following baseline, immersion media was either replaced by additional sterile saline, or terbutaline in sterile saline (1mM) to allow for pharmacological diffusion into the brain parenchyma and imaged for an additional 60 minutes. Baseline periods and saline treated animals controlled for microglial activation due to craniotomies and animals that had significant motility or morphological changes during the baseline were removed from the study.

#### Laser Ablation

Laser ablation injuries were created by running a point scan for 8s at 780nm using ∼75mW at the sample. The microglial injury response was imaged by collecting Z-stacks of 50-90μm every 5 minutes. For analysis, z-projections were all comprised of 10µm of the stack, encompassing the approximate center ablation core. The file was converted to an avi and subjected to analysis by a custom MATLAB script designed to calculate the movement of microglial processes towards the ablation core. Briefly, for each pixel at each time point the script generates a vector which estimates the magnitude and direction of motion of the pixel utilizing the Farneback method for estimating optic flow^15^. We then selected to only include vectors which were directed towards the ablation core to exclude motion not directed towards the injury core. We also filtered out vectors below 5 pixels of motion as these were identified as primarily noise in our imaging sessions. To quantify the generated vectors we summed the magnitudes of all the vectors at each time point and normalized this value to the total number of pixels in the image. The first time-point was excluded from all analysis as the script estimates the first time point’s movement by assuming a blank frame preceding the first time point. We then found the area under the curve for our injury response normalized magnitude as well as the maximum value of our normalized magnitude over the 1hr session.

### Code Availability

MATLAB code for motility and laser ablation analysis is available on request from the corresponding author.

### Histology

Whole brains were collected following transcardial perfusion and overnight post-fixation with paraformaldehyde (4%). Tissue was cryoprotected and coronal sections were cut on a freezing microtome (Microm; Global Medical Instrumentation, Ramsey, MN) at 50-μm thickness. Sections were processed free-floating at room temperature (RT). Briefly, sections were rinsed, and endogenous peroxidase activity and nonspecific binding blocked with a 10% BSA solution. Sections were then incubated in a primary antibody solution (24 h, 4°C, anti-Iba-1, 1:1,000, Wako #019-19741) followed by secondary antibody solution (4 h, RT, Alexa-Fluor 488, 1:500, Invitrogen), mounted and coverslipped. For examination of microglial ramification containing primary visual cortex were identified and imaged on a Zeiss LSM 510 confocal microscope (Carl Zeiss, Thornwood, NY). For each section, a 10um z-stack in the center of the tissue was collected with a z-step of 1 um at 40x magnifications. Analysis was performed offline in ImageJ. Z-stacks were smoothed and compressed into a single z-projection. For analysis of ramification, microglia whose entire process arbor was contained within the image were individually selected and cropped into a new image. Each image was thresholded to generate a binarized outline of the process arbor, filtered to remove artifacts and analyzed with an automated Sholl analysis plugin (kindly provided by the Anirvan Ghosh laboratory, UCSD).

### Statistics

Statistical comparisons were made between animal cohorts using Prism VI statistical analysis software. (GraphPad, La Jolla, CA). All n-values represent individual animals and are comparable to standard n values used in similar experiments in the literature. All values reported are the mean± s.e.m. For all analyses, α=0.05. Two-tailed unpaired, or paired t-tests and one-way or two-way ANOVAs with Bonferroni or Dunnett’s post hoc comparisons were used to compare cohorts where appropriate.

## Acknowledgements

We thank the University of Rochester Medical Center Flow Core for their expert training and services. We thank Cassandra Lamantia for assistance with animal management, John Olschowka and Kerry O’Banion for shared PCR resources, Fatima Rivera-Escalera for providing training on tissue preparation for microglial FACS, Anirvan Ghosh for the Sholl analysis ImageJ plugin, and Jianhua Cang for sharing Marlab code for OD analysis, and Brendan Whitelaw for assistance writing MATLAB scripts. This work was supported by the Nation Institutes of Health (NIH) grants R01 EY019277 (A.K.M), R21 NS099973 (A.K.M), F31 NS105249 (R.D.S), T32 NS007489 (R.D.S, G.O.S), F31 NS086241 (G.O.S), F32 EY028028 (G.O.S), National Science Foundation grant NSF 1557971 (A.K.M), Schmitt Program on Integrative Brain Research grant (G.O.S, R.P.D), University of Rochester Bilski-Mayer Fellowship (H.N.B), and University of Rochester Medical Center Summer Scholars Fellowship (K.A.L.).

## Author Contributions

R.D.S, G.O.S, and A.K.M conceived the project. R.D.S, G.O.S, R.P.D, H.N.B, and K.A.L. carried out experiments and data analysis. R.D.S carried out iOS imaging experiments and analysis in Cx3cr1CreERT/β2fl/fl, Cx3cr1CreERT, C57BL6J mice, and all *in vivo* two-photon experiments characterizing awake v. anesthetized mice, adrenergic pharmacological agents and imaging experiments utilizing Cx3cr1CreERT/β2fl/fl/Ai9, Cx3cr1CreERT/Ai9 mice. R.D.S also performed all FACS preparation on microglial specimens. G.O.S carried out iOS imaging experiments in C57BL6J mice and imaging experiments characterizing stress and circadian rhythms. G.O.S also performed experiments utilizing terbutaline. 4. R.P.D carried out stress experiments. H.N.B assisted in confirmation of Cx3cr1Cre/β_2_fl/fl excision. K.A.L carried out circadian morphology experiments. R.D.S, G.O.S, and A.K.M wrote the first draft of the manuscript. All authors contributed to the final version of the manuscript.

