## Supplemental Figures for "Noradrenergic signaling in wakeful states inhibits microglial surveillance and synaptic plasticity in the mouse visual cortex"

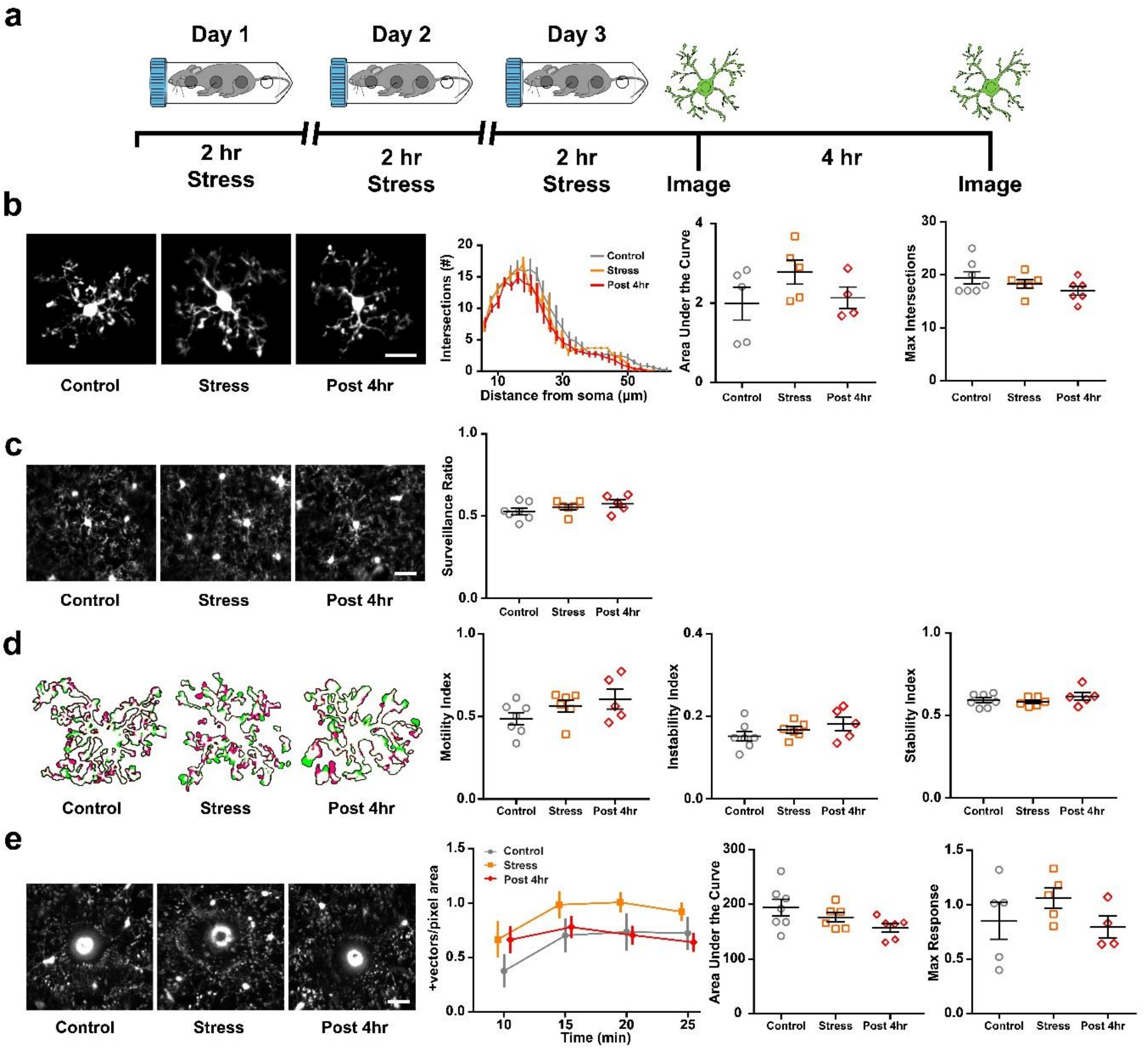


**Supplementary Figure 1| Acute stress does not affect microglial morphology, motility, or injury response.** (**a**) Schematic of the stress paradigm. Mice were restrained in a translucent tube under bright light for 2 hours daily for three days before imaging. Control mice were handled, but were returned to their cage. Microglia were imaged immediately following the last restraint period or 4 hrs later. (**b**) Left: example microglial morphologies for control, stressed or stressed +4hr mice (scale = 25µm). Left middle: sholl analysis demonstrating no difference in number of intersections as a function of distance from soma across all conditions. Right middle: area under curve (AUC) quantification showing no significant distance to overall process lengths. Right: Max number of intersections (maximal ramification) was also not significantly different between all conditions. (**c**) Left: example images of surveillance over a 30 minute period for control, stress and stress +4hrs (post 4hr) conditions (scale = 25µm). Middle: Area under curve (AUC) was not significantly different across all conditions. Right: The surveillance ratio was also not significantly different between all conditions. (**d**) Left: example overlays of microglial motility showing extension (green), retraction (magenta), and stable (white) areas over 5 minute time lapse. Left middle: no significant difference in motility index was found across all conditions. Right middle: no significant difference in instability index was found across all conditions. Right: no significant difference in stability index was found across all conditions. (**e**) Left: example microglial responses to laser ablation after 30 minutes in each condition (scale = 25μm). Left middle: no difference was observed for the response vectors as a function of time, though there was a trend for increased response in the stress condition. Right middle: area under curve (AUC) was not significantly different. Right: no significant difference was observed for the maximum response vector.

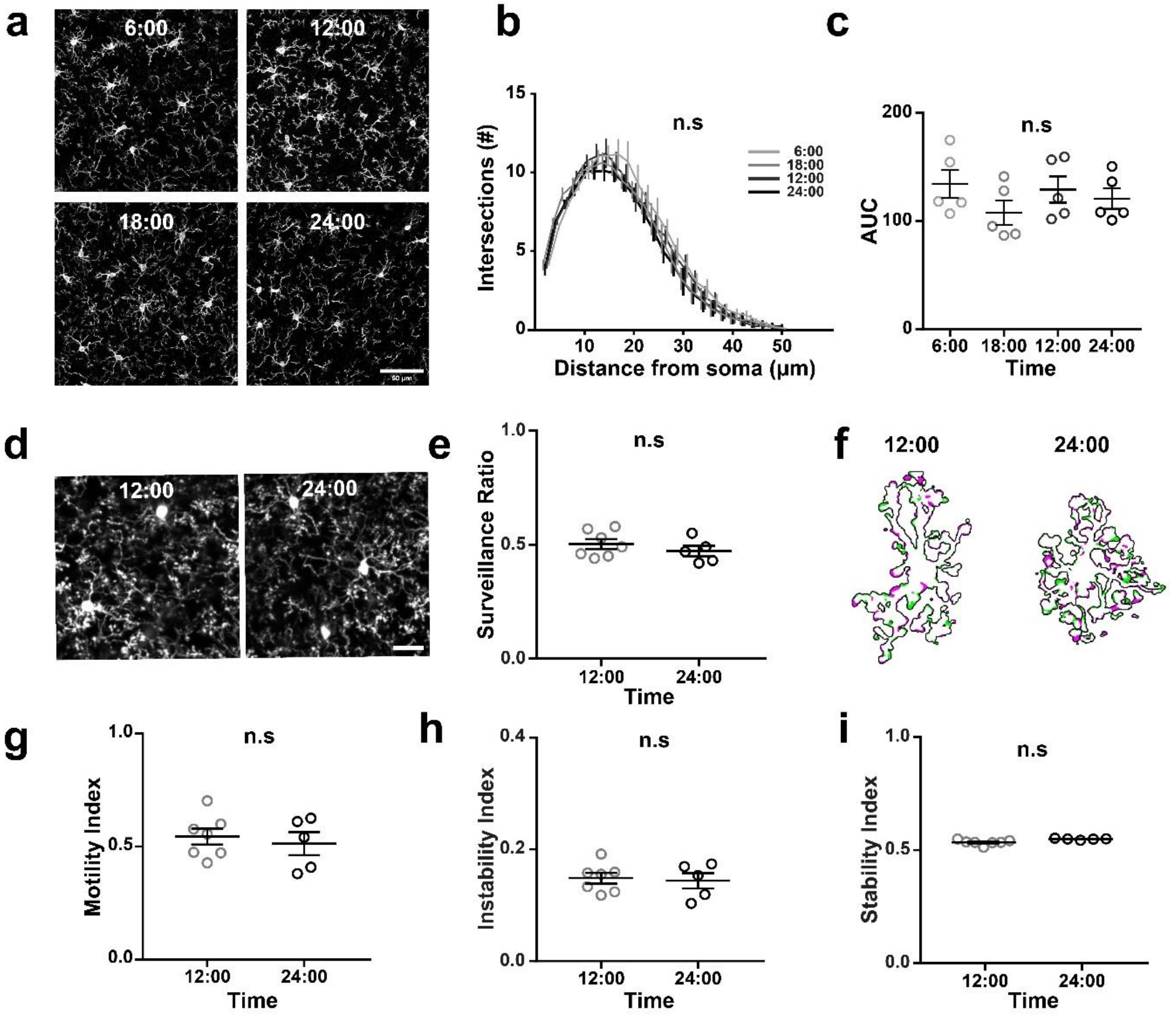


**Supplementary Figure 2| Circadian rhythm does not affect microglial morphology or motility dynamics.** (**a**) Example images of microglial morphology in fixed tissue stained for the microglial marker, Iba-1. Animals were sacrificed at four time points through the dark/light cycle corresponding to zeitgeber 6, 12, 18 and 24 (scale = 50µm). (**b**) Sholl analysis for microglia across at each timepoint. No difference was observed at any distance from the soma. (**c**) No difference was found for the total number of intersections (area under curve, AUC). (**d**) Example microglial process surveillance over 30 minutes imaged at zeitgeber 12 and 24 (scale = 25µm). (**e**) Quantification of surveillance ration showed no significant difference between either time point. (**f**) Example microglial time point overlays over 3 minutes showing extension (green), retraction (magenta), and stable areas (white). No difference was observed in motility index (**g**), instability index (**h**), or stability index (**i**).

**a**

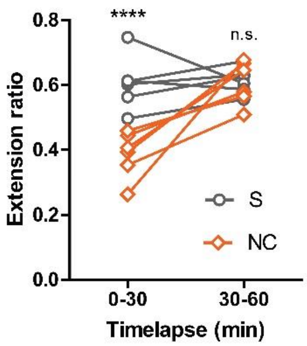


**Supplementary Figure 3| Clenbuterol transiently decreased microglial pseudopodia extension.** (**a**) In addition to the decrease in pseudopodia extension observed in clenbuterol-treated mice, we see a concomitant transient loss of pseudopodia extension (n=6 S, and n=6 NC, Two-way repeated measures ANOVA, P=0.0039, F(1,10)=13.91, Bonferroni post-tests for multiple comparisons, p<0.0001 s v. nc 0-30min).

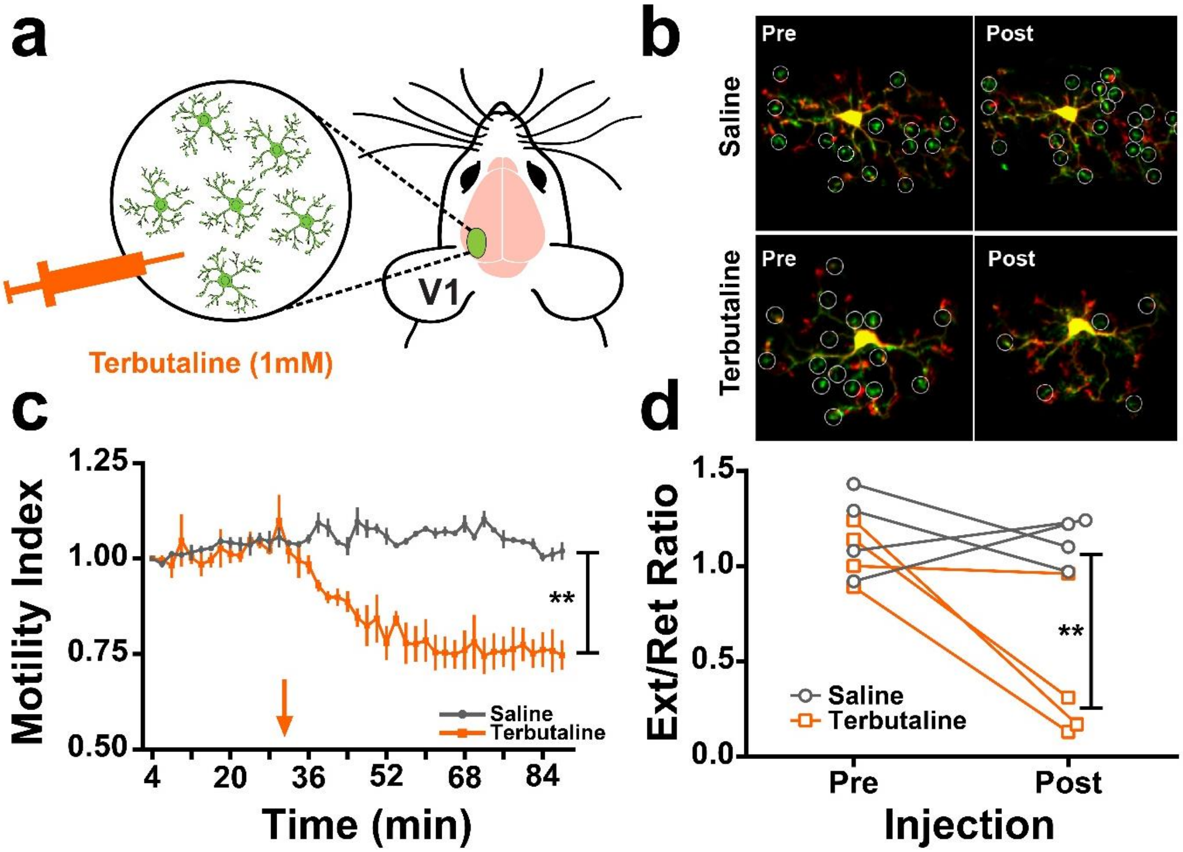


**Supplementary Figure 4| Direct terbutaline application reduces motility index and pseudopodia extension in microglia.** (**a**) Schematic of experimental setup. The β2-AR agonist, terbutaline (1mM), was applied directly to microglia (green) through an open cranial window in the visual cortex (V1) of anesthetized mice. (**b**) Example images of microglia before and after either saline administered controls (top) or terbutaline administered animals (bottom). Time lapse overlays correspond to the first 30 minutes of baseline, and the first 30 minutes following injection where extension (green), retraction (red), and stable areas (yellow) are represented. Circles highlight pseudopodia extensions. Note the decreased number of pseudopodia following terbutaline administration. (**c**) Motility index plotted as a function of time for the baseline period (time before red arrow), and after injection of saline (gray) or terbutaline (orange). Terbutaline causes a quick reduction in overall motility index. (**d**) Quantification of the total number of extension pseudopodia divided by the number of retraction pseudopodia (example shown in panel b). No difference was observed prior to saline (gray) or terbutaline (orange) administration (Pre, left). However, a significant reduction in pseudopodia extension is observed over the last half hour following terbutaline administration (Post, right). A ratio of 1.0 represents approximately equal number of extension and retraction pseudopodia.

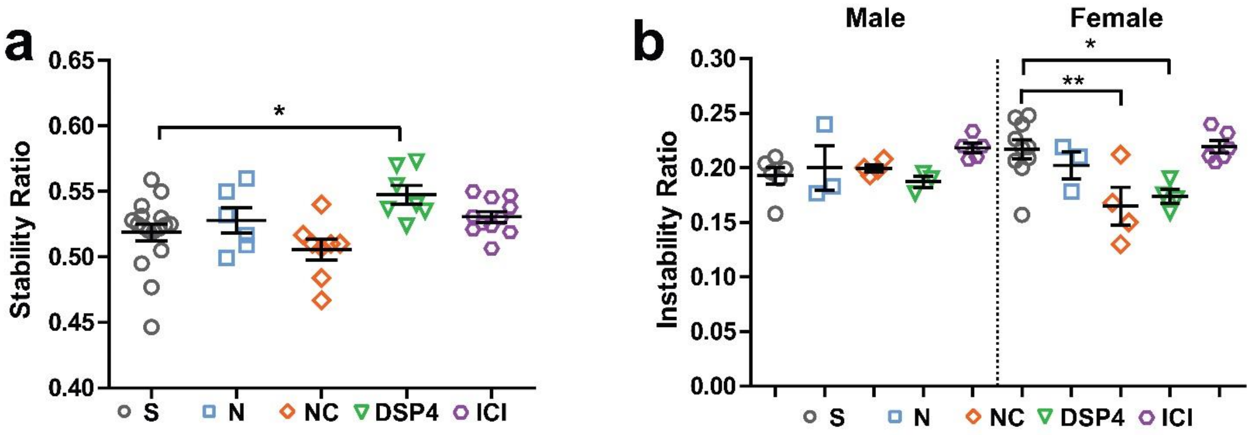


**Supplementary Figure 5| DSP4 stabilizes microglial processes while clenbuterol destabilizes microglial processes.** (**a**) Microglial process stability is increased in DSP4 treated mice (n=5-17 per group, One-way ANOVA, P=0.0080, F(4,48)=3.950, Dunnett's multiple comparisons tests, S v. DSP4 P<0.05). (**b**) Microglial instability is decreased in female clenbuterol and DSP4-treated mice (n=5-17 per group, One-way ANOVA, P=0.0448, F(4,38)=2.701, Dunnett's multiple comparisons tests, in females: S v. NC P<0.01 and S v. DSP4 P<0.05).

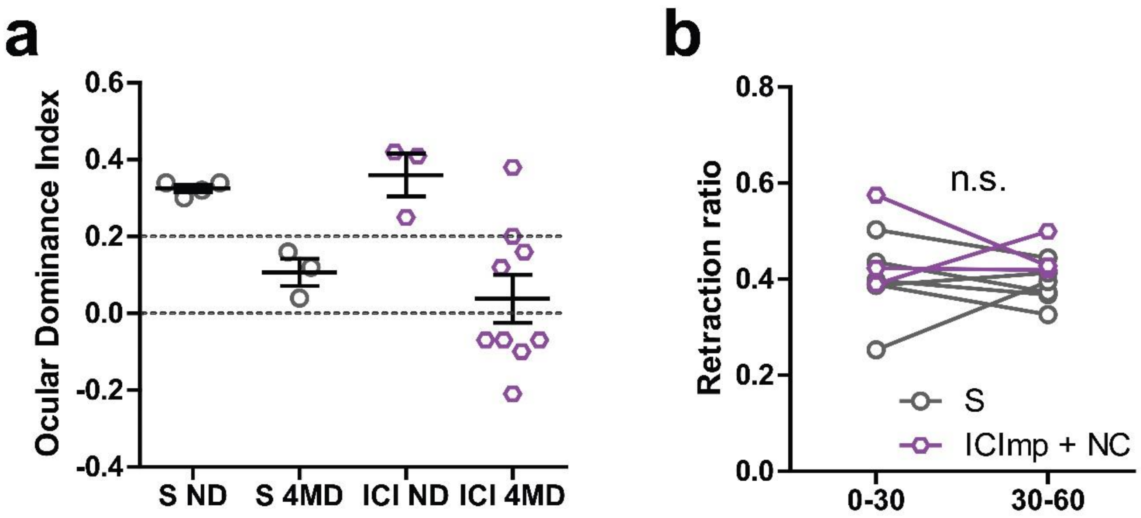


**Supplementary Figure 6| ICI 118,551 does not alter ocular dominance plasticity.** (a) Quantification of ocular dominance indices demonstrates robust shifts in saline and ICI 118,551 treated groups (n=3-9 per group, Two-way ANOVA, no significant interaction). (**b**) Imaging assay to confirm efficacy of mini-osmotic pump administration of ICI 118,551. We see ICI 118,551 given for 24hrs by mini-osmotic pump blocking the effects of clenbuterol *in vivo* (n=3-6 per group, Two-way ANOVA, no significant interaction).

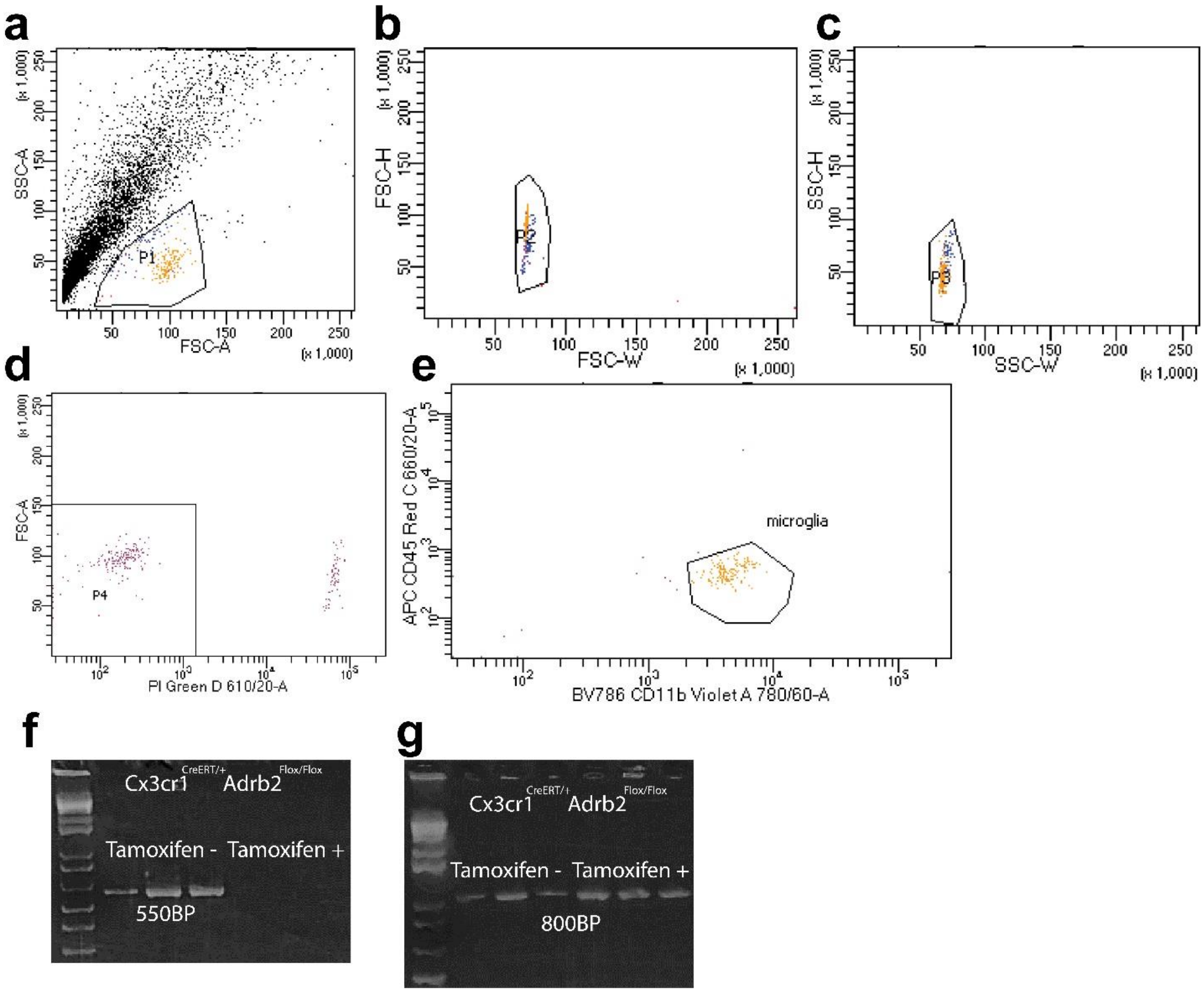


**Supplementary Figure 7| Cx3cr1-Cre$^{ERT}$ successfully drives excision of β$_2$ARs in microglia.** (**a-e**) Example FACS for microglia, microglia were sorted as PI negative (live), CD45$^{lo}$, and CD11b$^{Hi}$. (**f**) PCR for the presence of the β$_2$AR floxed allele, in mice given tamoxifen we no longer have the presence of the allele, suggesting excision (3 replicates per condition, replicates are individual animals). (**g**) PCR for the presence of the floxed excision product, we see recombination in both the untreated and tamoxifen-treated mice due to leakiness of the Cre (3 replicates per condition, replicates are individual animals).

### Supplementary Table 1

| Figure | Condition | # Male | # Female | Figure | Condition | # Male | # Female |
|---|---|---|---|---|---|---|---|
| **Fig 1a-j** | All | 4 | 5 | **Fig7e,f** | S ND | 5 | 5 |
| **Fig 1k,l** | All | ? | ? | | S 4MD | 5 | 4 |
| **Fig 2a-i** | All | 4 | 3 | | N ND | 1 | 3 |
| **Fig 3a-e** | All | 3 | 5 | | N 4MD | 4 | 1 |
| **Fig4a-c** | S | 3 | 3 | | NC ND | 2 | 4 |
| | NC | 3 | 3 | | NC 4MD | 3 | 7 |
| **Fig4 d-g** | S | 2 | 5 | | DSP4 ND | 1 | 5 |
| | N | 2 | 1 | | DSP4 4MD | 5 | 2 |
| | NC | 3 | 4 | **Fig7g,h** | S ND | 4 | 5 |
| **Fig4h-j** | S | 6 | 10 | | S 4MD | 8 | 7 |
| | N | 4 | 4 | | N ND | 2 | 3 |
| | NC | 4 | 4 | | N 4MD | 6 | 3 |
| | DSP4 | 3 | 4 | | NC ND | 3 | 2 |
| | ICI | 5 | 6 | | NC 4MD | 6 | 3 |
| **Fig5b,c,e,j** | NC | 3 | 3 | | DSP4 ND | 3 | 2 |
| **Fig5b,d,f,k** | ICI | 4 | 3 | | DSP4 4MD | 2 | 3 |
| **Fig5h** | NC | 3 | 3 | **SFig1** | | ? | ? |
| **Fig5i** | ICI | 3 | 3 | **SFig2** | | ? | ? |
| **Fig6 a-e** | S | 2 | 3 | **SFig3** | S | 3 | 3 |
| | N | 4 | 3 | | NC | 3 | 3 |
| | NC | 2 | 3 | **SFig4** | | ? | ? |
| | DPS4 | 2 | 4 | **SFig5** | S | 6 | 10 |
| | ICI | 2 | 4 | | N | 4 | 4 |
| **Fig7a,b** | S ND | 11 | 4 | | NC | 4 | 4 |
| | S 4MD | 8 | 6 | | DSP4 | 3 | 4 |
| | N ND | 5 | 1 | | ICI | 5 | 6 |
| | N 4MD | 4 | 2 | **SFig6a** | S ND | 2 | 1 |
| | NC ND | 2 | 4 | | S 4MD | 1 | 2 |
| | NC 4MD | 5 | 6 | | ICI ND | 2 | 1 |
| | DSP4 ND | 4 | 2 | | ICI 4MD | 7 | 2 |
| | DSP4 4MD | 2 | 6 | **SFig6b** | S | 3 | 3 |
| **Fig7c,d** | Cre Ai9 S | 3 | 1 | | ICI | 2 | 1 |
| | Cre Ai9 NC | 0 | 4 | **SFig7f,g** | CreB2 | 1 | 2 |
| | CreB2 Ai9 S | 2 | 3 | | CreB2+Tam | 0 | 3 |
| | CreB2 Ai9 NC | 4 | 1 | | | | |

**Supplemental video 1**

Time-lapse movie taken through a chronic cranial window showing the motility of V1 microglia in awake mice taken over 1 hour at 5-minute intervals (30μm z-stacks were compressed in each time point). Scale bar = 20μm

**Supplemental video 2**

Time-lapse movie showing the motility of V1 microglia in fentanyl cocktail anesthetized mice taken over 1 hour at 5-minute intervals (30μm z-stacks were compressed in each time point). Scale bar = 20μm

**Supplemental video 3**

Time-lapse movie taken through a chronic cranial window showing the motility of V1 microglia in dexmedetomidine anesthetized mice taken over 1 hour at 5-minute intervals (30μm z-stacks were compressed in each time point). Scale bar = 20μm

**Supplemental video 4**

Time-lapse movie taken through a chronic cranial window of V1 microglia in awake mice taken over 1 hour at 5 minute intervals after focal laser injury (10μm z-stacks were compressed at each time point). Scale bar = 20μm

**Supplemental video 5**

Time-lapse movie taken through a chronic cranial window of V1 microglia in fentanyl cocktail anesthetized mice taken over 1 hour at 5 minute intervals after focal laser injury (10μm z-stacks were compressed at each time point). Scale bar = 20μm

**Supplemental video 6**

Time-lapse movie taken through an acute craniotomy over V1 showing 30 minute baseline microglial motility at 5 minute intervals followed by the retraction of microglial pseudopodia with application of 1mM Clenbuterol for 30 minutes post-administration. Scale bar = 20μm.

**Supplemental video 7**

Time-lapse movie taken through a thin-skull preparation showing the motility of V1 microglia in saline dosed mice taken over 1 hour at 5-minute intervals (30μm z-stacks were compressed in each time point). Scale bar = 20μm

**Supplemental video 8**

Time-lapse movie taken through a thin-skull preparation showing the motility of V1 microglia in nadolol dosed mice taken over 1 hour at 5-minute intervals (30μm z-stacks were compressed in each time point). Scale bar = 20μm

**Supplemental video 9**

Time-lapse movie taken through a thin-skull preparation showing the motility of V1 microglia in nadolol/clenbuterol dosed mice taken over 1 hour at 5-minute intervals (30μm z-stacks were compressed in each time point). Scale bar = 20μm

**Supplemental video 10**

Time-lapse movie taken through a thin-skull preparation showing the motility of V1 microglia in DSP4-treated mice taken over 1 hour at 5-minute intervals (30μm z-stacks were compressed in each time point). Scale bar = 20μm

**Supplemental video 11**

Time-lapse movie taken through a thin-skull preparation showing the motility of V1 microglia in ICI-118,551-treated mice taken over 1 hour at 5-minute intervals (30μm z-stacks were compressed in each time point). Scale bar = 20μm

**Supplemental video 12**

Time-lapse movie taken through a chronic cranial window showing the motility of V1 microglia in awake ICI-118,551-treated mice taken over 1 hour at 5-minute intervals (30μm z-stacks were compressed in each time point). Scale bar = 20μm

**Supplemental video 13**

Time-lapse movie taken through a thin-skull preparation of V1 microglia in nadolol-treated mice taken over 1 hour at 5 minute intervals after focal laser injury (10μm z-stacks were compressed at each time point). Scale bar = 20μm

**Supplemental video 14**

Time-lapse movie taken through a thin-skull preparation of V1 microglia in nadolol/clenbuterol-treated mice taken over 1 hour at 5 minute intervals after focal laser injury (10μm z-stacks were compressed at each time point). Scale bar = 20μm
